# Five-layer systems analysis of *Leishmania* stage differentiation reveals an essential role for protein degradation in parasite development

**DOI:** 10.1101/2025.03.24.644963

**Authors:** Pascale Pescher, Thibaut Douché, Quentin Giai-Gianetto, Karen Druart, Julie Kovarova, Blaise Li, Thomas Cokelaer, K Shanmugha Rajan, Laura Piel, Céline Besse, Anne Boland, Jean-François Deleuze, Mariette Matondo, Michael P. Barrett, Shulamit Michaeli, Gerald F. Späth

## Abstract

Vector-borne, protist parasites have evolved complex developmental programs to adapt to very distinct host environments. How these important pathogens transition between insect and mammalian stages is only poorly understood. Here we investigate stage differentiation in the trypanosomatid parasite *Leishmania* that shows constitutive gene transcription, thus providing a unique model system to assess how development is governed by post-transcriptional mechanisms. Using a five-layer integrative systems analysis (from genome to metabolome), we examined hamster-isolated *Leishmania donovani* amastigotes and culture-derived, insect-stage promastigotes. This approach enabled us to rule out genomic adaptation as a key driver of parasite stage differentiation, while confirming the pivotal role of differential mRNA turnover in stage-specific gene expression. Assessing transcriptomic against proteomic expression changes uncovered an unexpectedly broad dynamic range of stage-regulated changes in protein abundance that only poorly correlated with mRNA levels. This discrepancy correlated with (i) alterations in snoRNA expression and changes in rRNA modification they guide suggesting stage-specific adaptation of the protein translation apparatus that can uncouple mRNA from protein abundancies, and (ii) differential protein degradation as revealed by quantitative proteomics of parasites treated with the proteasomal inhibitor lactacystin. Lactacystin treatment stalled the transition of spleen-derived amastigotes into promastigotes in culture, further underscoring the role of proteasomal activity in stage differentiation. Integration of our five-layer systems analysis established the first link between *Leishmania* development and the expression of co-regulated genetic networks encompassing mRNA turnover, protein translation, phosphorylation, and degradation. Our findings provide a powerful new resource for research programs that aim to dissect the emergent properties of regulatory networks and feedback loops underlying *Leishmania* stage differentiation, serving as a blueprint for other vector-borne pathogens that rely on disease-associated developmental transitions.

## INTRODUCTION

During their infectious cycle, many protist and fungal pathogens undergo complex developmental transitions that adapt their biology to different host environments, thus ensuring transmission, persistence or immune evasion. For example, stage differentiation of the malarial parasite *Plasmodium* spp is tightly controlled at transcriptional levels by the dynamic interplay between epigenetic regulators that remodel the chromatin landscape to accommodate stage-specific transcription factors, such as AP2 DNA-binding family members (1, 2). Likewise, the transition from fast-growing tachyzoites to slow-growing bradyzoites during mammalian *Toxoplasma* infection is regulated by the single master TF BFD1 (3). Unlike these apicomplexan parasites, stage-differentiation in the trypanosomatid pathogen *Leishmania* spp is not regulated at the level of transcriptional control, raising the question of how these parasites establish and maintain the various adaptive forms that develop inside their insect and mammalian hosts.

*Leishmania* spp are protist parasites of humans that represent a global public health problem causing a series of immunopathologies termed the leishmaniases (4–7). These parasites differentiate into various developmental forms that are adapted for extracellular proliferation inside the sand fly midgut (procyclic promastigotes), transmission from the vector to the mammalian host during uptake of a blood meal (metacyclic promastigotes), and intracellular proliferation inside fully acidified macrophage phagolysosomes (amastigotes) (8). Unlike other eukaryotes, *Leishmania* development is not controlled transcriptionally as protein-coding genes in these parasites lack individual promoters and are constitutively transcribed at all stages involving parasite-specific processes such as polycistronic transcription and trans-splicing (9, 10). Despite the largely constitutive gene transcription, different *Leishmania* stages are characterized by striking morphological, biochemical and metabolic changes (11–16). These are regulated by a plethora of post-transcriptional mechanisms, including differential mRNA turn over through the binding of various proteins and protein complexes to the 5’ and 3’ untranslated regions (UTRs) (17–22), that can trigger mRNA degradation via deadenylation and decapping (23, 24). Differential gene expression in *Leishmania* can further be regulated at the translational level, for example through (i) selective ribosome recruitment by of one of the six isoforms of the *Leishmania* cap-binding protein eIF4E that bind to the 5’ UTR, (ii) stabilizing the poly(A) tail via the recruitment of Poly(A) Binding Protein 1 (PABP1) to the 3’ UTR promoting translation, or (iii) the differential expression of ribosomal proteins that can increase the landscape of specialized ribosomes (25–34). Finally, differential gene expression in *Leishmania* can also be controlled at the level of protein stability, through stage-specific expression of proteasomal components, autophagy-related genes (ATGs) or lysosomal proteases (35–41).

While each level of gene expression control has been studied individually in different *Leishmania* species, how these different types of regulation are integrated during *Leishmania* stage differentiation remains to be elucidated. Here, we approached this open question by applying a 5-layer, systems level analysis on the two major life cycle stages of *Leishmania donovani*, i.e. hamster-isolated, *bona-fide* amastigotes and derived promastigotes. Our analysis draws a complex picture of the *Leishmania* differentiation process that emerges from complex, co-regulated genetic networks involving mRNA turnover, protein translation, protein phosphorylation and protein degradation. Our data reveal *Leishmania* as an interesting model system to analyze phenotypic adaptation in the absence of transcriptional control and to dissociate the role of complex regulatory networks in the development of stable, disease-causing life cycle stages in eukaryotic pathogens.

## MATERIAL & METHODS

### Ethics statement

Work on animals was performed in compliance with French and European regulations on care and protection of laboratory animals (EC Directive 2010/63, French Law 2013-118, February 6th, 2013). All animal experiments were approved by the Ethics Committee and the Animal welfare body of Institut Pasteur and by the Ministère de l’Enseignement Supérieur, de la Recherche et de l’Innovation (projects n°#19683 and #240013).

### Animals

Twenty-five female Golden Syrian hamsters (*Mesocricetus auratus* RjHan:AURA, weighting between 50 – 60 g) were purchased from Janvier Laboratories. All animals were handled under specific, pathogen-free conditions in biohazard level 3 animal facilities (A3) accredited by the French Ministry of Agriculture for performing experiments on live rodents (agreement A75-15-01).

### Parasites and culture

*Leishmania donovani* strain 1S2D (MHOM/SD/62/1S-CL2D) was obtained from Henry Murray, Weill Cornell Medical College, New York, USA and maintained by serial passages in hamsters. Anesthetized hamsters were inoculated by intra-cardiac injection of 5×10^7^ amastigotes purified from hamster infected spleens as described previously (42). The weight of the animals was recorded over time, and the animals were euthanized by CO_2_ asphyxiation before they reached the end-point of infection represented by a 20% loss of body weight. Amastigotes were then recovered from the infected hamster spleens and used for nucleic acid and protein extractions, or differentiated into promastigotes at 26°C in M199 complete medium (M199, 10% FBS, 25 mM HEPES; 100 µM adenine, 2 mM L-glutamine, 10 µg/ml folic acid, 13.7 µM hemin, 4.2 mM NaHCO_3_, 1xRPMI1640 vitamins, 8 µM 6-biopterin, 100 units penicillin and 100 µg/ml streptomycin, pH 7.4). Promastigotes, derived from splenic amastigotes, were collected after 2 *in vitro* passages for nucleic acid and protein extractions.

### Experimental design

Strains issued from independent experimental evolution assays are identified by number, e.g. amastigote ama1 and promastigote pro1 are the parasites prepared from hamster H1 (see Figures 1A and S1 for details). For comparative analyses, DNA, RNA, proteins and metabolites were extracted from different splenic ama and their matching pro parasites, the latter being collected for extraction during exponential growth phase.

**Figure 1:**
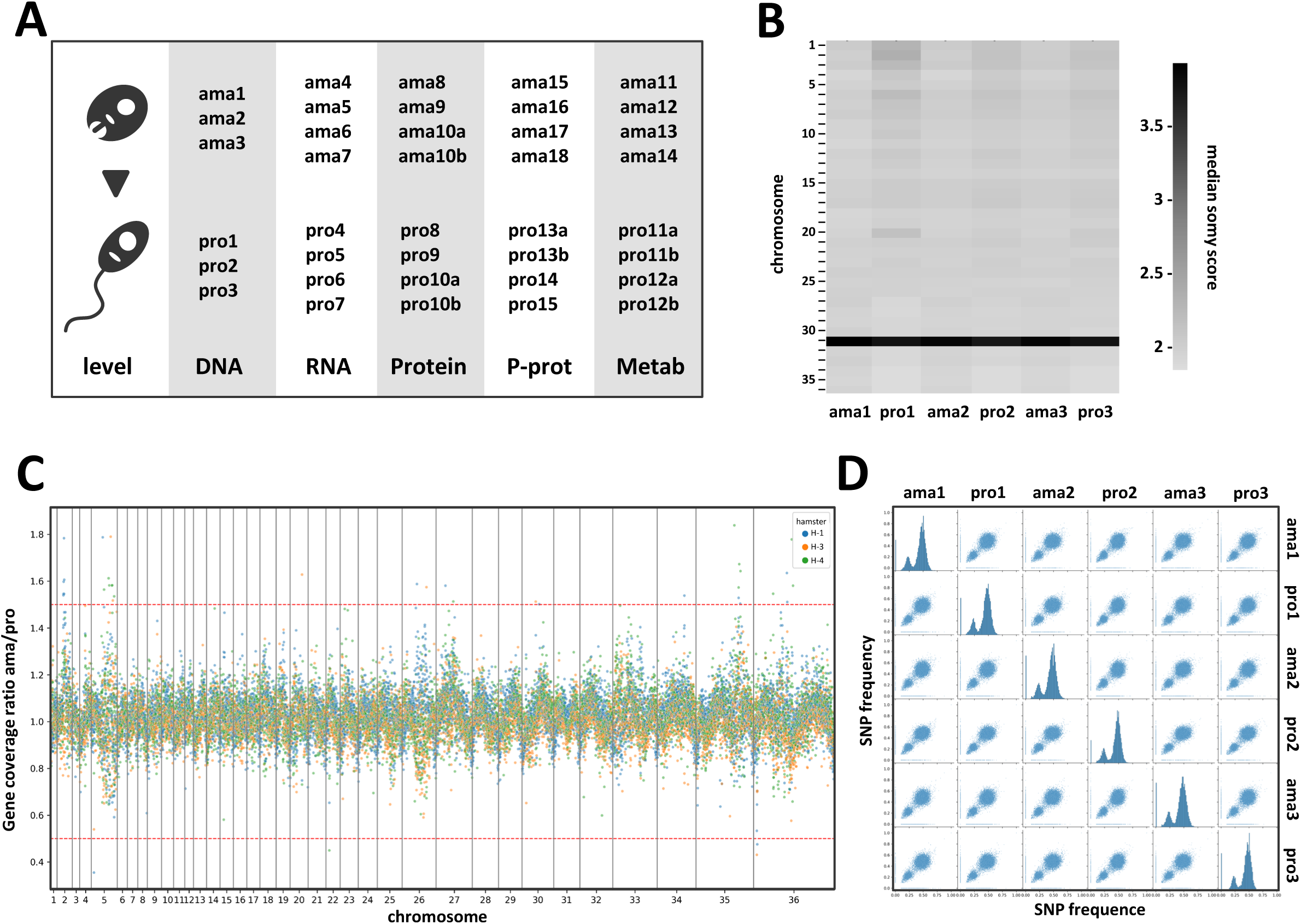
Comparative genomic analysis of *L. donovani* lesion-isolated amastigotes and derived promastigotes. (A) Schematic overview of the parasite samples used for the various systems level analyses presented in this study. A more detailed overview is shown in Figure S1. (B - D) Genomic analyses. (B) Heatmap showing the somy score of three individual differentiation experiments using hamster-isolated amastigotes (ama1, ama2 and ama3) and their corresponding, culture-derived promastigotes analysed at passage 2 (pro1, pro2 and pro3). Samples and chromosomes are indicated on the x- and y-axis, respectively. The somy scores were calculated as described in Material and Methods and correlate to the grey level according to the shown legend. (C) Ratio plot showing the gene coverage ratio ama versus pro (y-axis) for all genes across the 36 chromosomes (x-axis) as calculated based on median read depth normalized by the somy score. Each color represents one individual differentiation experiment using ama isolates from individual hamsters. The red dotted lines indicate the ratio values corresponding to 1.5 (upper line) and 0.5 (lower line), indicating respectively gain or loss of one gene copy in the ama samples. Fluctuations between 0.5 and 1.5 were not considered significant. (D) Correlation plots representing the individual (histograms on the diagonal) and pairwise (off-diagonal scatterplots) distributions of SNP frequencies for the indicated samples. Of note, only SNPs with a frequency above 10% were plotted. Due to bottle neck events, some low frequency SNPs appear to be unique in one or the other stage as they are either ‘lost’ (filtered out) or ‘gained’ (passing the 10% cutoff) between the ama and pro samples. The absence of SNPs at 100% is explained by the heterozygous nature of the Ld1S genome.

### Parasite differentiation and lactacystin treatment

Spleen-derived amastigotes were incubated at 2×10^7^/ml and 26°C in presence or absence of 10 µM of lactacystin (L6785, Sigma) for 3, 6, 18 and 48h in M199 complete medium. At each time point, parasites were collected to control for the presence of paraflagellar rod protein 2 (PFR2) as promastigote differentiation marker. After one *in vitro* passage, amastigote-derived promastigotes were incubated at a concentration of 2×10^7^ parasites per ml in presence or absence of 10 µM of lactacystin for 18h. Parasite viability was assessed by FACS analysis after 18h of treatment by propidium iodide (1µg/ml) or YO-PRO-1 (0.2 μM) (# Y3603, Invitrogen) staining as readout using untreated and PFA-treated parasites as controls. Parasites were collected from 25 ml of culture after 18h of treatment for quantitative, label-free proteomics analyses.

### Microscopy

Differential interference contrast (DIC) microscopy was applied on hamster-derived amastigotes during *in vitro* differentiation to promastigotes using an Axioplan 2 imaging microscope using a 63x oil immersion objective, the AxioVision Rel.4.8 software and an AxioCam MRm camera (Carl Zeiss). Alternatively, microscopic images of amastigotes in presence or absence of the lactacystin inhibitor were obtained using the EVOS FL microscope (life technologies) at a 20x magnification.

### Western blot analysis

Protein extracts were obtained from parasites after 3, 6, 18 and 48h of differentiation from amastigotes to promastigotes in presence or absence of lactacystin. Extracts were loaded on 4-12% SDS-PAGE (NuPage 4-12% Bis-Tris gel NP0321BOX, Invitrogen), separated by electrophoresis and stained with Coomassie blue or transferred onto PVDF membrane. The presence of PFR2 and α-tubulin was revealed using the anti-PFR2 antibody (kindly given by Philippe Bastin), an anti- α-tubulin antibody (clone B-5-1-2, Sigma), the corresponding HRP-conjugated secondary antibodies (Invitrogen) and the SuperSignal West Pico PLUS kit (Thermoscientific). Protein extracts from lactacystin-treated or untreated amastigotes and promastigotes were labelled with Cy5, loaded on 4-12% SDS-PAGE (NuPage 4-12% Bis-Tris gel NP0321BOX, Invitrogen), separated by electrophoresis and transferred onto PVDF membrane. The presence of ubiquitinated proteins was revealed using the mouse anti-ubiquitin monoclonal antibody P4D1 (Abcam), the HRP-conjugated anti-mouse secondary antibody (Invitrogen) and the ECL Prime western blotting detection reagent (Cytiva). Images were acquired using the ImageQuant 800 (Cytiva).

### Genome sequencing and data analysis

DNA was prepared from splenic amastigotes and promastigotes at exponential culture phase (three biological replicates). Parasites were centrifuged at 1,600 x *g* (pro) or 2,000 x *g* (ama) for 10 min at room temperature. Approximately 1×10^8^ promastigotes from logarithmic growth phase were re-suspended in 200 µl PBS and genomic DNA was purified using DNeasy Blood and Tissue kit from Qiagen and RNase A, according to the manufacturer’s instructions. DNA concentrations were measured in duplicate by fluorescence using a Molecular Device fluorescence plate reader (Quant-IT kits, Thermo Fischer Scientific). The quality of DNA was controlled by determining the DNA Integrity Number (DIN) analyzing 20 ng of DNA on a TapeStation 4200 (Agilent). One µg of genomic DNA was used to prepare a library for whole genome sequencing on an automated platform, using the Illumina “TruSeq DNA PCR-Free Library Preparation Kit”, according to the manufacturer’s instructions. After normalization and quality control, qualified libraries were sequenced on a HiSeqX5 platform from Illumina (Illumina Inc., CA, USA) at the Centre National de Recherche en Génétique Humaine (CEA, Evry, France), generating paired-ended, 150-bp reads. Sequence quality parameters were assessed throughout the sequencing run. Standard bioinformatics analysis of sequencing data was based on the Illumina pipeline to generate a FASTQ file for each sample.

Genomic DNA reads were aligned to the *L. donovani* Ld1S reference genome (https://www.ncbi.nlm.nih.gov/bioproject/PRJNA396645, GCA_002243465.1) with BWA mem (version 0.7.12) with the flag -M to mark shorter split hits as secondary. Samtools fixmate, sort, and index (version 1.3) were used to process the alignment files and turn them into bam format (43). RealignerTargetCreator and IndelRealigner from the GATK suite were run to homogenize indels (44). Eventually, PCR and optical duplicates were labeled with Picard MarkDuplicates (version 1.94(1484)) (https://broadinstitute.github.io/picard/) using the option “VALIDATION_STRINGENCY=LENIENT”. For each read alignment file, Samtools view (version 1.3) and BEDTools genomecov (version 2.25.0) were used to measure the sequencing depth of each nucleotide (45). Samtools was run with options “-q 50 -F 1028” to discard reads with a low map quality score or potential duplicates, while BEDTools genomecov was run with options “-d -split” to compute the coverage of each nucleotide. The coverage of each nucleotide was divided by the median genomic coverage. This normalization is done to account for library size differences. The chromosome sequencing coverage was used to evaluate aneuploidy between EP.1 and LP.1 samples. Then for each sample and for each chromosome, the median sequencing coverage was computed for contiguous windows of 2,500 bases. As previously published (46), the stably disomic chromosome 36 was used to normalize chromosome read depth and to estimate chromosome polysomy levels in each sample. Gene counts were produced using featureCounts (version 1.4.6-p3 (47)) with these parameters: -s 0 -t gene - g gene_id and were normalized according to the median-ratio method.

For the generation of the chromosome median somy score heatmap, mean coverage in 300 bp bins as generated by the GIP pipeline (48) were used to compute somy scores per chromosome by first normalizing bin scores for a sample by their median across all genome (to obtain comparable values between samples), multiplying by two (to scale somy values to the default diploid state assumed for most of the genome), and then taking the median across bins belonging to a given chromosome (see also https://gip.readthedocs.io/en/latest/giptools/karyotype.html). These chromosome scores were then displayed on a sample x chromosome heatmap, excluding the “maxi” chromosome. This was done using the Pandas 1.4.2 (49, 50) Pandas-dev/pandas: Pandas 1.2.4, https://zenodo.org/records/4681666 (50)), Matplotlib 3.5.1 (51) and Seaborn 0.11.2 (52) Python libraries.

For the assessment of gene coverage ratios, normalized mean coverages per gene are reported by GIP (version 1.1.0 (48)), as the mean coverage of the gene divided by the median coverage of the chromosome containing the gene. For a given gene, and a given amastigote - promastigote pair, a gene coverage ratio was computed adding 0.1 to the gene coverages of the individual samples, in order to avoid zero division errors. The genes were sorted by genomic coordinate and assigned successive integers to create a “genomic index”. Gene coverage ratios were then plotted along this genomic index, using the same Python libraries as above.

For the SNP analysis, alternative (Alt) allele frequencies for SNPs were retrieved from the output of the SNV giptools module (provided along with the GIP pipeline). Euclidean distances between samples in the SNP allele frequencies space were computed and used to build a dendrogram using the scipy.cluster.hierarchy Python module (SciPy version: 1.8.0). The clustering was obtained using the UPGMA method.

### RNA-seq analysis

Total RNA was extracted from splenic amastigotes and promastigotes in exponential culture phase (4 biological replicates). Amastigotes and promastigotes were centrifuged respectively at 2,000 x *g* or 1,600 x *g* for 10 min at 4°C, and re-suspended in the lysis buffer supplied with the Qiagen RNeasy Plus kit. The samples were stored at -80°C and RNA extractions were performed according to the manufacturer’s instructions, including a DNase treatment. RNA integrity was validated using the Agilent Bioanalyzer. DNase-treated RNA extracts were used for library preparation. Libraries were constructed using an Illumina Stranded mRNA Prep (Illumina, USA) following the supplier’s recommendations. RNA sequencing was performed at the Biomics Center (Institut Pasteur, Paris, France) on the Illumina NextSeq 2000 platform for a target of 40M paired-end reads per sample.

The RNA-seq analysis was performed using the Sequana RNA-seq pipeline (version 0.20.1, https://github.com/sequana/sequana_rnaseq) from the Seauqnq project (53). The pipeline workflow manager is Snakemake 7.3.2 (54). The pipeline was executed with default parameters and bioinformatics software are available as reproducible containers from the Damona project (https://github.com/cokelaer/damona). Reads were mapped to the *Leishmania donovani* reference using bowtie2 2.4.5 (55). Gene-level quantification was conducted using FeatureCounts 2.0.1 (47) assigning reads to genomic features based on the gene annotation and gene_id attribute, while accounting for strand-specificity information. Differential expression analysis was performed using DESeq2 v1.24.0 (56) scripts available in the Sequana library. Statistical testing identified differentially expressed genes by comparing amastigote and promastigote sample groups, with significance determined using Benjamini-Hochberg adjusted p-values (false discovery rate FDR < 0.05).

To specifically assess the differential expression of snoRNA, RNA-seq libraries from promastigote and amastigote stages were mapped to the Ld1S genome, and snoRNA read counts were obtained using bedtools multiBamCov. For snoRNAs with multiple copies, read counts were summed per sample to generate a single expression value per snoRNA family. DESeq2 was then used to perform differential expression analysis between amastigote and promastigote stages based on the summed snoRNA counts.

### Label-free, quantitative total proteome analyses

Untreated amastigotes (ama and ama-18h) and promastigotes (pro), and lactacystin-treated parasites (ama lacta and pro lacta) were recovered from four replicates (Figure 1A and S1) and washed three times with cold PBS at 2,000 x *g* or 1,600 x *g* for 10 min at 4°C. Parasite lysates were prepared in eFASP lysis buffer (4% SDS / 0.2 % DCA / 50mM TCEP / 50 mM ammonium bicarbonate buffer pH 8) (61). For detailed protocol see supplementary data file. MS scans were acquired at a resolution of 70,000 and MS/MS scans (fixed first mass 100 m/z) at a resolution of 17,500. The AGC target and maximum injection time for the survey scans and the MS/MS scans were set to 3 x 10^6^ for 20 ms and 10^6^ for 60 ms, respectively. An automatic selection of the 10 most intense precursor ions was activated (Top 10) with a 40 s dynamic exclusion. The isolation window was set to 1.6 m/z and normalized collision energy fixed to 28 for HCD fragmentation. We used a minimum AGC target of 10^4^ corresponding to an intensity threshold of 1.7 x 10^5^. Unassigned precursor ion charge states as well as 1, 7, 8 and >8 charged states were rejected and match was disabled.

### Label-free quantitative phosphoproteome analyses

Amastigote (ama) and promastigote (pro) parasites from four biological replicates were recovered and washed three times by centrifugation in cold M199 at respectively 2,000 x *g* or 1,600 x *g* for 10 min at 4°C. Samples were incubated for 10 min at 4°C in lysis buffer (1ml per 1.5×10^9^ parasites) consisting of 8 M urea, 50 mM Tris, supplemented with a protease inhibitor cocktail (cOmplete from Roche) and a phosphatase inhibitor cocktail (PhosStop from Roche). Following sonication for 5 min using a sequence of 10 s pulse and 20 s pause, the lysates were centrifuged 15 min at 14,000 x *g* and 4°C, and the supernatant was collected and stored at −80°C until use. Proteins were quantified by RC DC protein assay (Bio-Rad) and adjusted to 1.3 µg.µl^-1^ in lysis buffer. For detailed protocol see supplementary data file. Phosphopeptide enrichment was carried out as described in (62) and detailed in supplementary data. All analyses were performed on a Q Exactive HF Mass Spectrometer (Thermo Fisher Scientific) coupled with a Proxeon EASY-nLC 1000 (Thermo Fisher Scientific) as detailed in supplementary data. Details for the proteomics and phosphoproteomics analyses following the data acquisition are available in supplementary data. Results are presented in table 6 and 10.

Phosphopeptides were selected for Gene Ontology enrichment analyses if (i) they were exclusively identified in one of the two stages, (ii) they showed significant, stage-specific changes in phosphopeptide abundance (fold change ≥ 2, adj. p-value < 0.01) even if the corresponding protein was not detected in the total proteome analysis, and (iii) they showed a significant increase in relative phosphorylation normalized to protein abundance (p-value < 0.05) as calculated by the ratio ‘change in phosphopeptide abundance’ vs ‘change in protein abundance’ using a cut off of fold changes ≥ 2 and adj. p-value < 0.01 for both analyses (see table 11).

### Metabolomic analysis

The sample extraction was performed as described previously (63). Briefly, 10^8^ cells were used per each 200 μL sample (4 replicates/stage). Firstly, cells were rapidly quenched in a dry ice/ethanol bath to 4°C, then centrifuged, washed with 1 x PBS, and resuspended in the extraction solvent (chloroform:methanol:water, 1:3:1). After vigorous shaking at 4°C for 1 h, extracts were centrifuged (16,000 x g, 4°C, 10 min) and the supernatants collected and stored at −80°C until the analysis. LC-MS analyses were performed using separation on 150 x 4.6 mm, 5 mm ZIC-pHILIC (Merck) on UltiMate 3000 RSLC (Thermo Scientific) followed by mass detection on an Orbitrap Exactive mass spectrometer (Thermo Fisher) at Glasgow Polyomics. Analyses were performed in positive and negative polarity switching mode, using 10 μl injection volume and a flow rate of 300 μl/min over 26 min on the column maintained at 30°C, as follows: 0 to 20 min 20%-to 80% solution A, 15 to 17 min 95% solution A, 17 to 26 min 20% solution A where solution A is 20 mM ammonium carbonate in water and solution B is acetonitrile). The samples were run alongside 249 authentic standards at 10 μM each. Mass spectrometry data was processed using Mzmatch (64) and Ideom (63) software. Unique signals were extracted using the centwave algorithm and matched across biological replicates based on mass-to-charge ratio and retention time. These grouped peaks were then filtered based on relative standard deviation and combined into a single file. The combined sets were then filtered on signal-to-noise score, minimum intensity and minimum detections. The final peak set was then gap-filled and converted to text for use Putative metabolite identification corresponds for the most part to Metabolite Standards Initiative (MSI) level 2 (mass only, and thus only considered a tentative annotation), whereas metabolites matching in retention time to an included standard correspond to level 1 (considered likely an accurate annotation). Peaks having an area with root squared deviation across pooled samples > 50% were excluded, as were those with a retention time < 4 min (due to poor resolution).

### Localizing rRNA modification on *Leishmania* cryo-EM ribosome structure

The cryo-EM atomic model of *Leishmania major* 80S ribosomes bound to mRNA and all three tRNAs (PDB: 8RXH) was used to project rRNA modifications of the ribosome. Figures were generated using UCSF Chimera-X software (65).

### Systems-level analyses

A lists of *L. donovani* GO terms (66) was built in house. The Biological Networks Gene Ontology tool (BiNGO) plugin of the Cytoscape software package (version 3.8.2) was used to map and visualize functional enrichments in each data set based on the GO hierarchy. A Benjamini & Hochberg false discovery rate with a significance level of 0.05 was applied. Cluster efficiency represents the percentage of genes for a given GO term compared to the total number of genes with any GO annotation in the considered set of genes. Enrichment score corresponds to the percentage of genes for a given GO term compared to all the genes sharing the same GO term in the genome. Word Cloud of GO enrichment analysis limited to GO Slim terms and a threshold of p-value < 0.05 for the category Biological Process was performed on TriTrypdb (https://tritrypdb.org) using the *L. donovani* LdBPK orthologs for all genes that showed statistically significant expression changes (p-value < 0.01) at both RNA or protein levels.

For the phosphoproteomic work, phosphopeptides were selected for Gene Ontology enrichment analyses if (i) they were exclusively identified in one of the two stages, (ii) they showed significant, stage-specific changes in phosphopeptide abundance (fold change ≥ 2, adj. p-value < 0.01) even if the corresponding protein was not detected in the total proteome analysis, and (iii) they showed a significant increase in relative phosphorylation normalized to protein abundance (p-value < 0.05) as calculated by the ratio ‘change in phosphopeptide abundance’ vs ‘change in protein abundance’ using a cut off of fold changes ≥ 2 and adj. p-value < 0.01 for both analyses (see table 11).

Functional enrichment networks were built with the STRING plugin of the Cytoscape software package (version 3.8.2) using the *L. infantum* orthologs, and the full STRING network with a confidence score cutoff of 0.4. A false discovery rate with a significance level of 0.05 was applied. The “ClusterProfiler” package (version 4.2.2) of R was used for KEGG (Kyoto encyclopedia of genes and genomes) gene set enrichment analysis (GSEA) and for data mapping on metabolic pathways available in the KEGG database. Results were visualized using the “pathview” packages (version 1.34.0).

### Data availability

The mass spectrometry proteomics data were deposited to the ProteomeXchange Consortium via the PRIDE partner repository with the dataset identifiers PXD035697 (phosphoproteome data) and PXD035698 (proteome upon lactacystin treatment) (67). The DNAseq data have been deposited in NCBI’s Gene Expression Omnibus (68) and are accessible in the BioProject PRJNA1231373 (https://www.ncbi.nlm.nih.gov/bioproject/PRJNA1231373). The RNAseq data have been deposited in ArrayExpress (EMBL-EBI) (69) under the accession number E-MTAB-16528. The mass spectrometry metabolomic data are available on Figshare at https://doi.org/10.6084/m9.figshare.31027849. The scripts for GIP and giptools are available at https://github.com/susefranssen/Global_genome_diversity_Ldonovani_complex.

## RESULTS

### *Leishmania* stage differentiation occurs independently from changes in gene dosage

We carried out an in-depth, systems analysis of independent *L. donovani* amastigote preparations isolated from infected hamster spleens and their culture-derived promastigotes at *in vitro* passage 2, with the aim of revealing mechanisms controlling stage-specific differential expression across five quantifiable information levels, by comparing the genome, transcriptome, proteome, metabolome and phospho-proteome of the two life-cycle stages (Figure 1A and Figure S1). We first investigated the possible role of genome instability in stage-differentiation, given the very well documented propensity of *L. donovani* to establish and tolerate chromosome and gene copy number variations in culture and during sand fly or hamster infection (46, 66, 70).

Applying our computational pipeline termed ‘GIP’ (48) to the genome sequences of three independent pairs of amastigotes and promastigotes confirmed the largely disomic state of the *Leishmania* genome at both stages, except for the known constitutive tetrasomy of chr 31 (Figure 1B and Figure S2A, Table 1). We did not detect any major changes at karyotypic levels following promastigote differentiation, confirming our previous observations that stage differentiation occurs independent of karyotypic adaptation (46, 70). Nevertheless, a clear pattern emerged for promastigotes that showed a slight but reproducible increase in median coverage for chr 1, 2, 3, 4, 6 and 20, suggesting the emergence of mosaic aneuploidy for these chromosomes early during culture adaptation. However, these minor signals are not predictive for the karyotypic changes characteristic of culture-adapted Ld1S parasites (i.e. chr 5, 26, 33, (46, 70)). Likewise, differentiation of promastigotes was not associated with any major gene copy number variation at early passage 2 (Figure 1C, Table 2). Even though a number of genes showed increased (e.g. Ld1S_300477100 encoding an intraflagellar transport (IFT) protein) or decreased (e.g. Ld1S_220235700 encoding a hypothetical protein) read depth in individual pro strains, none of these changes converged across all pro samples. In the absence of convergent selection, it is impossible to distinguish if these gene CNVs provide some strain-specific advantage or are merely the result of random genetic drift. Plotting the normalized read depth ratios between amastigotes and promastigotes revealed an intriguing, undulating pattern that was highly reproducible across the three independent experiments and has been previously associated with nascent DNA produced during the replicative S-phase of the cell cycle (71–73). Finally, differentiation did not affect the SNP frequency distribution (Figure S2B, Table 3), which resulted in a largely diagonal pattern when plotting amastigote against promastigote SNP frequencies (Figure 1D).

In conclusion, unlike long-term culture adaptation, which drives important karyotypic changes (46, 74) that can affect transcript- and protein abundance levels as demonstrated by a recent 4-layer systems analysis (75) *in vitro* differentiation of splenic amastigotes into promastigotes was not associated with significant changes in chromosome or gene copy number, ruling out gene dosage effects as the source of stage-specific expression differences. Thus, *L. donovani* promastigote differentiation *in vitro* is independent of genomic adaptation, confirming our previous results obtained with sand fly-derived, promastigote parasites (66, 76).

### Comparative RNA-seq analysis reveals stage-specific, co-regulated gene clusters that are controlled at post-transcriptional levels

We next analyzed stage-specific expression changes applying RNA-seq analyses on four amastigote and derived promastigote strains (Figure 2 and Figure S3). In contrast to the relatively stable genome infrastructure maintained between stages, we observed substantial, stage-specific differences in transcript abundance. From 10402 detected transcripts in both stages, 6478 showed statistically significant differences (adj. p-value < 0.01, base mean reads ≥ 10 reads), including 1234 and 1027 transcripts with a 2-fold or higher increase in abundance in amastigotes and promastigotes, respectively (Table 4). Considering the lack of promoter-driven expression control of individual, protein coding genes in *Leishmania* and the absence of stage-specific gene dosage effects (see Figure 1), the observed differences in amastigote and promastigote transcript abundances are most likely attributable to stage-specific mRNA turnover. Interestingly, non-coding (nc) RNAs account for 64.7% of the differentially expressed transcripts at the amastigote stage (Figures 2A).

**Figure 2:**
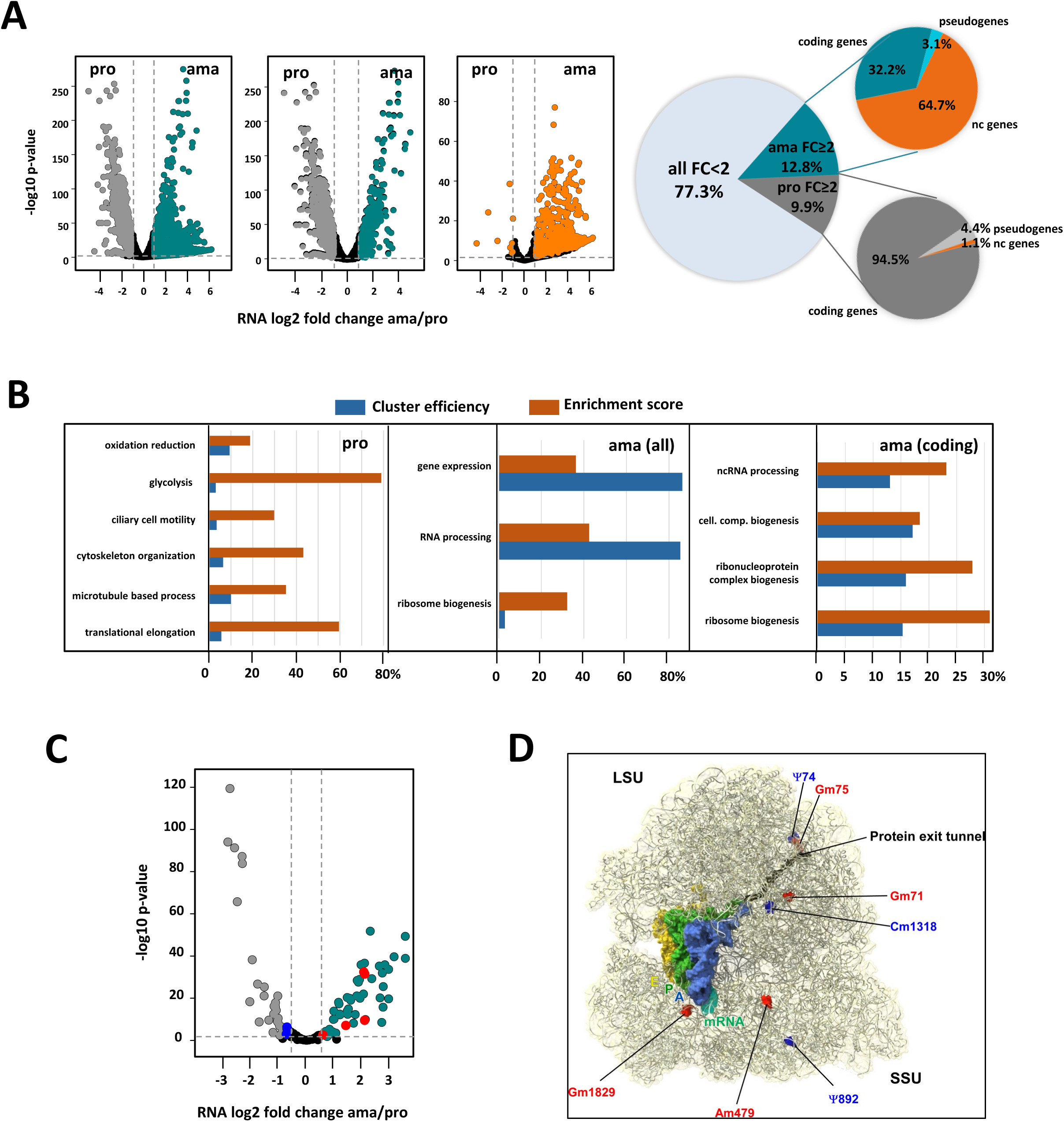
Stage-specific transcript profiling. (A) Volcano plots (left panels) showing differential transcripts abundance between ama and pro samples as assessed by RNA-seq analysis. The dotted lines indicate fold change (FC) = 2 (vertical line) and p-value = 0.01 (horizontal line). Transcripts with FC < 2 or adjusted p-value > 0.01 are represented by black dots. Transcripts with significant increased abundance FC ≥ 2 and adjusted p-value < 0.01 in ama and pro are indicated respectively in dark cyan and dark grey for all transcripts (left panel) or transcripts of only coding genes (middle panel). Transcripts for non-coding genes with significant increased abundance FC ≥ 2 and adjusted p-value < 0.01 in ama and pro are indicated in the right volcano plot (orange dots). The pie chart (right panel) depicts the percentage of transcripts for each of the categories. (B) GO term enrichment analysis of these transcripts for the category ‘biological process’ was performed, and ‘cluster efficiency’ (blue) and ‘enrichment score’ (orange) were plotted for pro (left panel) and ama datasets (all transcripts, middle panel; transcripts for coding genes, right panel). The graphs show some of the most significantly enriched GO terms associated with a Benjamini and Hochberg (BH) p-value < 0.05 that were identified using the BiNGO plug-in in Cytoscape. (C) Volcano plot corresponding to the differential expression of snoRNA between pro and ama stages. Cyan dots (ama) and grey dots (pro) represent signals with FC ≥ 2 and adj. p-value < 0.05. Black dots represent non-significant changes. (D) Localization of differentially modified rRNA sites on the *Leishmania* ribosome. The RNA modifications are indicated as space filling in the *Leishmania* cryo-EM structure (PDB: 8RXH). The identity of the RNA modification is defined by the label (Ψ, pseudouridine; Gm and Cm refer to guanine and cytosine methylation, respectively; numbers indicate residues). Upregulated and downregulated RNA modifications in amastigotes according to Rajan et al (34) are indicated in red and blue, respectively.

Analyzing the stage-specific RNA-seq data sets identified clusters of co-regulated transcripts that share a common annotation (Figure 2B, Table 4). In promastigotes, we observed increased abundance for 66 transcripts annotated for the interconnected GO terms ‘microtubule based process’, ‘cytoskeleton organization’ and ‘ciliary cell motility’, confirming the post-transcriptional co-regulation of flagellar biosynthesis as previously shown in *L. mexicana* (77) (Figure 2B left panel, Table 4, sheet G). Other co-regulated, functional gene sets are defined by the GO terms ‘oxidation reduction’ (46 transcripts), ‘glycolysis’ (15 transcripts), ‘post-transcriptional regulation of gene expression’ (46 transcripts) and ‘translational elongation’ (28 transcripts), further supporting translational control as a regulatory step in stage differentiation.

In amastigotes, a major co-regulated gene cluster is represented by 77 amastin transcripts encoded on 5 different chromosomes that show increased abundance in amastigotes, confirming the stage-specific, post-transcriptional regulation of this gene family as previously described (19, 78, 79) (Table 4, sheet C, E and F). Another functional gene cluster showing increased abundance in amastigotes was associated with protein translation, with GO enrichment observed for the terms ‘ribosome biogenesis’ (26 transcripts), ‘ncRNA processing’ (22 transcripts), including 17 transcripts annotated for the GO term ‘rRNA processing’, and 6 transcripts for ‘tRNA processing’ (Figure 2B, middle and right panel, Table 4, sheet E and F).

These data suggest ribosome and epi-transcriptomic regulation as possible central components in *Leishmania* stage differentiation. Indeed, analyzing poly-A transcriptome libraries of amastigotes and promastigotes we observed significant differences in stage-specific expression of 110 pre-snoRNAs (Table 5, Figure 2C). These data fit with the expression of a series of methyl- and pseudouridyl-transferases at this stage (see Table 6, sheets A and E). Stage-specific expression of some of these snoRNAs correlated with stage-specific changes previously observed in Pseudouridine and 2’-O-Methylation rRNA modifications (34). The positions of these modifications and the corresponding snoRNAs known to affect ribosome translational activity are indicated on the high resolution cryo-EM structure of the *Leishmania* ribosome (Figure 2D).

In conclusion, our RNAseq data reveal a series of functionally related, stage-specific gene clusters whose mRNA abundance is regulated in a coordinated manner at post-transcriptional level. Whether this co-regulation is governed by stage-specific mRNA decay mechanisms acting on 5’ and 3’ UTRs or by stage-specific mRNA stabilization via RNA-binding proteins or mRNA modification remains to be established.

### Quantitative proteome analysis reveals co-regulated gene clusters that are independent of stage-specific mRNA turnover

We next employed quantitative proteomic analysis to assess how the different co-regulated gene clusters observed in our RNAseq analyses translate into stage-specific proteomes. Label-free quantitative proteomic analysis of 4 replicates of amastigotes and derived promastigotes identified over 4000 proteins, including 1987 differentially expressed proteins (adjusted p-value < 0.01), and 1534 that were exclusively detected in either ama or pro (Figure 3A left panel, Table 6). Considering the 822 proteins that are specific or more abundant in amastigotes (fold change ≥ 2), functional enrichment was mainly observed for the GO terms ‘oxidation reduction’ (61 proteins), ‘vesicle mediated transport’ (23 proteins) and ‘phosphorylation’ (41 proteins) (Figure 3A, middle panel, Table 6, sheets B and D). Considering the 1637 proteins that are specific or more abundant in promastigotes (fold change ≥ 2), functional enrichment was observed for the GO terms ‘post-transcriptional regulation of gene expression’ (71), ‘translation’ (69 proteins), ‘ribosome biogenesis’ (21 proteins) and ‘ciliary cell motility’ (42 proteins) (Figure 3A, right panel, Table 6, sheet C and E).

**Figure 3:**
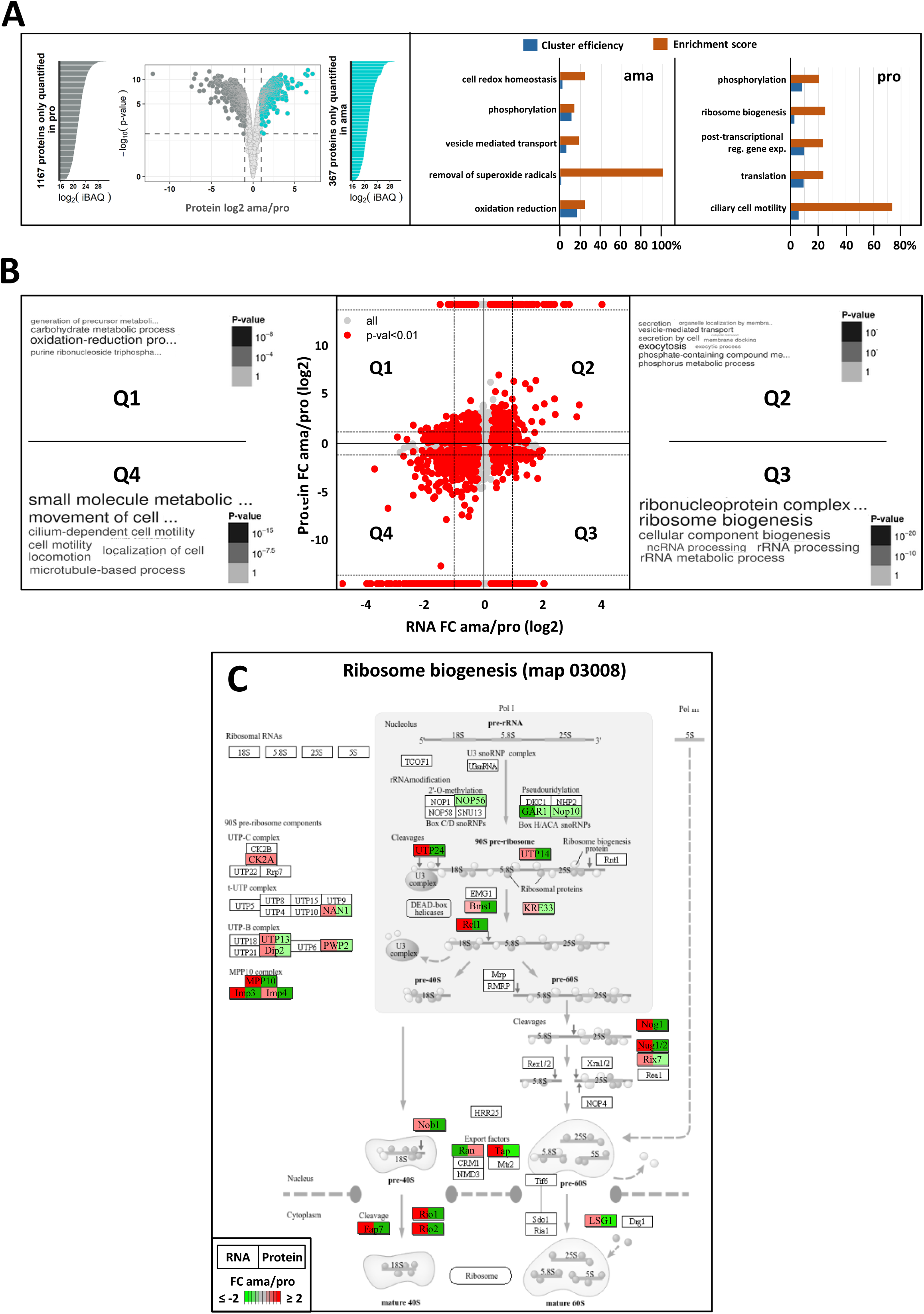
Systems analysis of the stage-specific transcriptome and proteome. (A) Volcano plot showing differential protein abundance between four ama and pro samples as assessed by label-free, quantitative proteomics analysis. The dotted lines indicate fold change (FC) = 2 (vertical line) and false discovery rate (FDR) = 0.01 (horizontal line). Proteins with FC < 2 or FDR > 0.01 are represented by light grey dots. Proteins that were reproducibly detected in only one of the two stages (considered unique) are represented by the lateral histograms plotting their relative abundance according the iBAQ value. These proteins together with proteins showing differential abundance of FC ≥ 2 and adjusted p-value < 0.01 are indicated for ama in dark cyan and for pro in dark grey (see Table 6). GO term enrichment analysis for the category ‘biological process’ was performed, and the resulting values for ‘cluster efficiency’ (blue) and ‘enrichment score’ (orange) were plotted for ama (middle panel) and pro (right panel). The graphs show the main GO terms identified using the BiNGO plug-in in Cytoscape and associated with a BH p-value < 0.05. (B) Double ratio plot showing the log2 fold change between ama and pro in transcript (x-axis) and protein (y-axis) abundances. Red dots correspond to changes in both transcript and protein abundance defined either by significant differential abundance at both levels (p-value < 0.01), or by significant RNA changes (p-value < 0.01) corresponding to proteins detected in only one of the two stages. The dotted lines indicate fold changes (FC) = 2. Word cloud enrichment performed with the *L. donovani* LdBPK orthologs is presented for each of the 4 quadrants, Q1 to Q4 and a detailed GO enrichment analysis is given in Table 7. The font size is proportional to the number of genes per GO term and the grey scale refers to the p-value calculated for each GO term. (C) KEGG map for ribosome biogenesis. Each gene associated with expression changes indicated in Q1 to Q4 (see Figure 2C) and annotated with the GO term ‘ribosome biogenesis’ has been projected on the ribosome biogenesis KEGG map. Each gene is represented by a square with the left segment showing fold changes between ama and pro at transcript and the right segment showing fold changes at the protein level, with the differential abundance indicated by the color and its intensity (red, increase; green, decrease). Only genes quantified in both transcriptome and proteome analyses and associated with an adjusted p-value < 0.01 were considered.

The metabolomic analysis of four independent amastigote and promastigote samples (Figure S4A) supported our proteomic data. Promastigotes showed increased amino acid biosynthetic activity that matched the changes in protein abundance observed for this pathway, while the amastigote metabolome indicates a shift from glycolytic to TCA cycle-dependent energy production that was previously described (12, 14, 80) (Figures S4B-D), with a notable increase in metabolites involved in butanoate metabolism indicative of a shift between carbohydrate use as a primary energy source to utilization of fatty acid oxidation. The amastigote expression profile and metabolome further indicate the stage-specific production of the storage-carbohydrate mannogen, thus corroborating previous observations (81) (Figure S4E).

Surprisingly, only few enriched GO terms matched between transcriptomics and proteomics datasets, suggesting additional layers of regulation between the transcriptional and translational levels of gene expression. To investigate this aspect on a genome-wide level, we next plotted the simple numerical values we obtained when calculating the ratios of expression changes between ama and pro observed in the transcriptomics and proteomics datasets (Figure 3B, Table 7). Only expression changes were considered that either showed statistically significant differential abundance at both RNA and protein levels (p < 0.01), or showed significant RNA changes (p < 0.01) with the corresponding protein being detected in only one of the two stages. These latter proteins are identified by signals that were arbitrarily placed at the upper (detected in ama) or the lower (detected in pro) parts of the graph. Whether these proteins just escape detection due to low expression or are truly not expressed remains to be established. The double ratio analysis of the resulting 2349 gene products defined four quadrants: Quadrants 2 (Q2) and 4 (Q4) correspond to 430 and 1042 genes (Table 7, sheet E and I), respectively, for which stage-specific expression changes correlate between protein and RNA levels, suggesting that the abundance of these proteins may be mainly regulated by mRNA turn-over. Q2 (mRNA and protein up in ama) includes the GO terms ‘vesicle mediated transport’ (17 genes) and ‘lipid metabolic process’ (21 genes) (Table 7, sheet F), while Q4 (mRNA and protein up in pro) includes the GO term ‘cell motility’ (41 genes) (Table 7, sheet J).

Counter-correlations between RNA and protein abundances (p-value < 0.01) were observed for 463 and 414 genes, respectively falling into quadrants Q1 (RNA down and protein up in ama) and Q3 (RNA down and protein up in pro) (Figure 3B, Table 7, sheets C and G). The most important difference was observed for 35 genes annotated for the GO term ‘ribosome biogenesis’ (Figure 3C, Table 7, sheet H). While this GO term defined a positively regulated gene cluster at mRNA level in amastigotes (see Figure 2B), the same genes show reduced expression at this stage at protein level (Figure 3C). This lack of correlation may be explained by stage-specific changes in mRNA translation efficiency.

In conclusion, while many stage-specific changes of protein abundance are strictly regulated by similar changes at mRNA level, others show discrepancies in stage-specific abundances between transcript and protein levels, revealing regulatory mechanisms that act independent of mRNA stability, for example at the level of translational control or differential protein stability.

### Stage-specific protein turn-over is required for parasite differentiation

The discrepancies we observed in a sub-set of genes between changes in mRNA and protein abundance during the amastigote-to-promastigote transition (see Figure 3B) primed us to investigate the role of proteasomal protein degradation in *Leishmania* differentiation. We chose the highly specific and irreversible proteasome inhibitor lactacystin (82, 83) over the typanosomatid-specific, reversible drug candidate LXE408 (84) as the latter’s potent cytotoxicity can confound direct effects on protein turnover with secondary consequences of cell death, limiting its utility for dissecting proteasome function in living parasites. Comparative, quantitative proteomics analyses were performed with (i) spleen-derived amastigotes (ama), (ii) amastigotes after the first 18h of differentiation in the absence (control, ama-18h) and the presence of latacystin (ama-lacta), and (iii) promastigotes in culture without (pro) and after 18h of treatment with lactacystin (pro-lacta) (Table 8, Figure 4A and S5).

**Figure 4:**
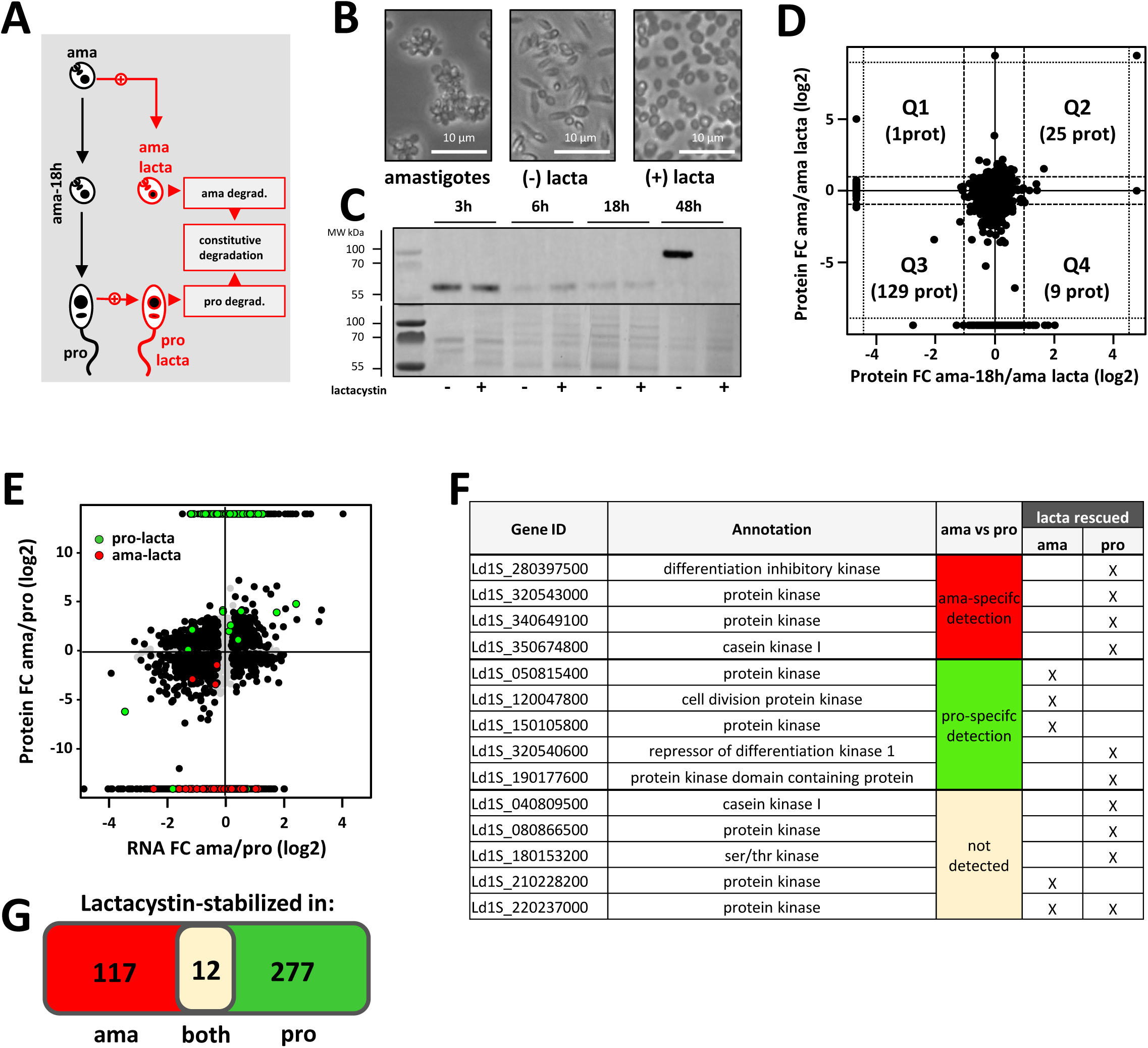
Analysis of stage-specific proteasomal protein turn-over. (A) Overview of experimental workflow. To assess early and late steps of differentiation (left part of the graph in black), comparative, quantitative proteomics analyses were performed on tissue-isolated amastigotes (ama), amastigotes at 18h into the transition to promastigotes in culture (ama-18h) and fully differentiated promastigotes after 1 passage *in vitro* (pro). To gain insight into stage-specific and constitutive protein degradation (right part of the graph in red), amastigotes and promastigotes were treated with the irreversible inhibitor lactacystin (ama-lacta and pro-lacta). (B) Microscopic images of amastigotes at 18h in absence of lactacystin treatment (amastigotes, left), after 48h in absence (-, middle) or presence of the inhibitor (+, right). (C) Western blot analysis. Protein extracts were obtained from 2×10^6^ parasites after 3, 6, 18 and 48h of differentiation from amastigotes to promastigotes in presence (+) or absence (−) of lactacystin. Extracts were separated by electrophoresis, transferred onto PVDF membrane and the presence of PFR2 was revealed using an anti-PFR2 antibody (upper panel). A Coomassie blue stain of the same gel is shown as loading control (lower panel). (D) Double ratio plot comparing the log2 fold changes in protein abundance between ama-18h/ama-lacta (x-axis) and ama/ama-lacta (y-axis). Dashed lines indicate FC = 2. (E) Projection of the proteins stabilized in presence of the inhibitor in ama (ama-lacta, orange) and pro (pro-lacta, pink) onto the double ratio plot shown in Figure 3B. Note that ama-specific proteins are rescued from degradation in pro-lacta, while pro-specific proteins are rescued from degradation in ama-lacta. (F) List of the protein kinases stabilized after lactacystin treatment in pro or ama (columns lacta rescued) and either not detected in the total proteome (column ama vs pro, grey cell) or specifically quantified at the ama (red cell) or the pro (green cell) stage. (G) Venn diagram showing the number of proteins specifically stabilized in ama or pro or stabilized in both stages after lactacystin treatment.

This data set allowed us important insight into processes of early differentiation, revealing increased abundance in ama-18h compared to ama for 694 proteins (fold change ≥ 2, adjusted p-value < 0.01) (Table 8b, sheet B) with GO enrichment observed for the terms ‘ribosome biogenesis’, ‘RNA methylation’, ‘rRNA processing’, ‘gene expression’, and ’post-transcriptional regulation of gene expression’, the latter including 6 genes encoding for pseudouridine transferases known to modify ribosomal (r) RNA (Figures S5A and S6) (Table 8b, sheets D to H). These data further sustain a role of epitranscriptomic regulation and stage-adapted ribosomes as key processes that may initiate *Leishmania* promastigote differentiation, at least in culture (31, 34, 66).

Lactacystin treatment further uncovered an essential regulatory role of proteasomal protein degradation in *L. donovani* development, which builds on previous published observations (85). While the presence of the inhibitor neither affected parasite viability (Figure S7A) nor morphology (data not shown and Figure S7B) or promastigote growth (as previously shown by Silva-Jardim et al (85)), lactacystin treatment abrogated the amastigote-to-promastigote developmental transition as judged by the persistence of an amastigote-like morphology (i.e. oval cell shape, retracted flagellum), and the absence of PFR2 expression characteristic of promastigotes (Figures 4B and C). In contrast, untreated amastigotes spontaneously converted to flagellate promastigotes after 48h in culture (Figure S7D). Proteasome inhibition blocked amastigote-to-promastigote differentiation, without inducing rapid global accumulation of ubiquitinated proteins (Figure S7C, upper panel) consistent with a quiescent-like state and low basal ubiquitin–proteasome system activity in amastigotes. After 18 h, ubiquitination levels remained similar to untreated cells, indicating that protein turnover and ubiquitin accumulation are primarily driven by developmental remodeling rather than acute proteasome inhibition. In promastigotes, the lack of detectable change (Fig. S7C, lower panel) may also reflect high basal ubiquitination, engagement of compensatory pathways such as autophagy, and/or only partial proteasome inhibition.

Comparing the proteomics signatures of lactacystin-treated samples and untreated controls (Figure S5) allowed us to reveal the proteasomal targets, whose rescue correlated with the observed abrogation of differentiation. This analysis is complicated by the fact that amastigotes will not only respond to the 18h treatment but also initiate their differentiation into promastigotes during this period, at least in the untreated control sample. To distinguish between changes caused by lactacysin treatment and changes due to the differentiation process, we plotted the ratio of ama/ama-lacta against the ratio ama-18h/ama-lacta (Figure 4D), which defined one quadrant (Q3) representing 129 proteins whose increased abundance in ama-lacta was exclusively due to rescue from degradation (Table 9, sheet A). Applying STRING and GO analyses on these proteins, some of the most significantly enrichment terms included ‘protein kinase’, ‘cellular response to stimulus’, and ‘cell cycle’ (Figure S8 lower panel, Table 9, sheet B and G).

Proteomics analysis of lactacystin-treated promastigotes (pro-lacta) revealed the rescue of 289 proteins from degradation when compared to untreated control (pro) (Table 9, sheet C). Comparison of rescued proteins across both ama and pro stages revealed a series of intriguing findings: First, most of these proteins were rescued in a stage-specific manner, confirming differential protein degradation as an important regulatory component in the development and maintenance of *Leishmania* life cycle stages as suggested by previous reports (37, 86–92). Second, just like the proteins rescued from proteasomal degradation in ama-lacta, the pro-lacta data set too showed GO enrichment for the terms ‘protein kinase’ and ‘cellular response to stimulus’, involving distinct sets of proteins (Figure S8, upper panel) (Table 9, sheets D, E and G). Thus, *Leishmania* differentiation correlates with the expression of complex signaling networks that are established in a stage-specific manner. Third, when projecting the lactacystin-rescued proteins to the established, stage-specific proteomes (see Figure 3), an intriguing pattern emerged: the majority of proteins that were rescued in ama-lacta turned out to be part of the promastigote proteome data set (Figure 4E). *Vice versa,* many proteins rescued in pro-lacta were part of the amastigote-specific proteome data set (Table 9, sheets H and I), a pattern that affected 14 protein kinases (Figure 4F). Thus, these proteins seem to be expressed constitutively across both stages, while their steady-state stage-specific abundance is regulated by proteasomal degradation. Surprisingly, 12 proteins (including the protein kinase Ld1S_220237000) were rescued from degradation in both ama-lacta and pro-lacta but were never identified in the proteomes of the corresponding untreated controls, suggesting that they are constitutively degraded in both stages under our experimental conditions (Figure 4G, Table 9, sheet F).

In conclusion, our results confirm the important role of protein degradation in regulating the *L. donovani* amastigote and promastigote proteomes and identify protein kinases as key targets of stage-specific proteasomal activities.

### Quantitative phosphoproteome analysis reveals *Leishmania* biological networks associated with stage differentiation

The investigations above associated the expression of protein kinases (and their degradation) to *Leishmania* stage differentiation (Figure 4F, S8 and S9), thus confirming previous reports on the importance of phosphotransferase activities in trypanosomatid development (93–103). Differential turnover of these important signaling proteins by stage-specific proteasomal activities likely represents a regulatory switch controlling parasite development. To further investigate how this putative regulatory switch operates, we screened for stage-specific protein kinase substrates applying label-free, quantitative phospho-proteomic analyses on three independent amastigote and promastigote samples. We identified a total of 2079 phospho-peptides in amastigotes and 7095 phospho-peptides in promastigotes, suggesting a three-fold increase in global protein phosphorylation in insect-stage parasites (Table 10). However, increased TiO_2_ phosphopeptide enrichment in a given sample may be due either to increased substrate expression or increased *de novo* phosphorylation (Figure 5A and S10). To distinguish between these two possibilities we assessed stage-specific relative phosphorylation changes normalized to protein abundance (Figure 5B).

**Figure 5:**
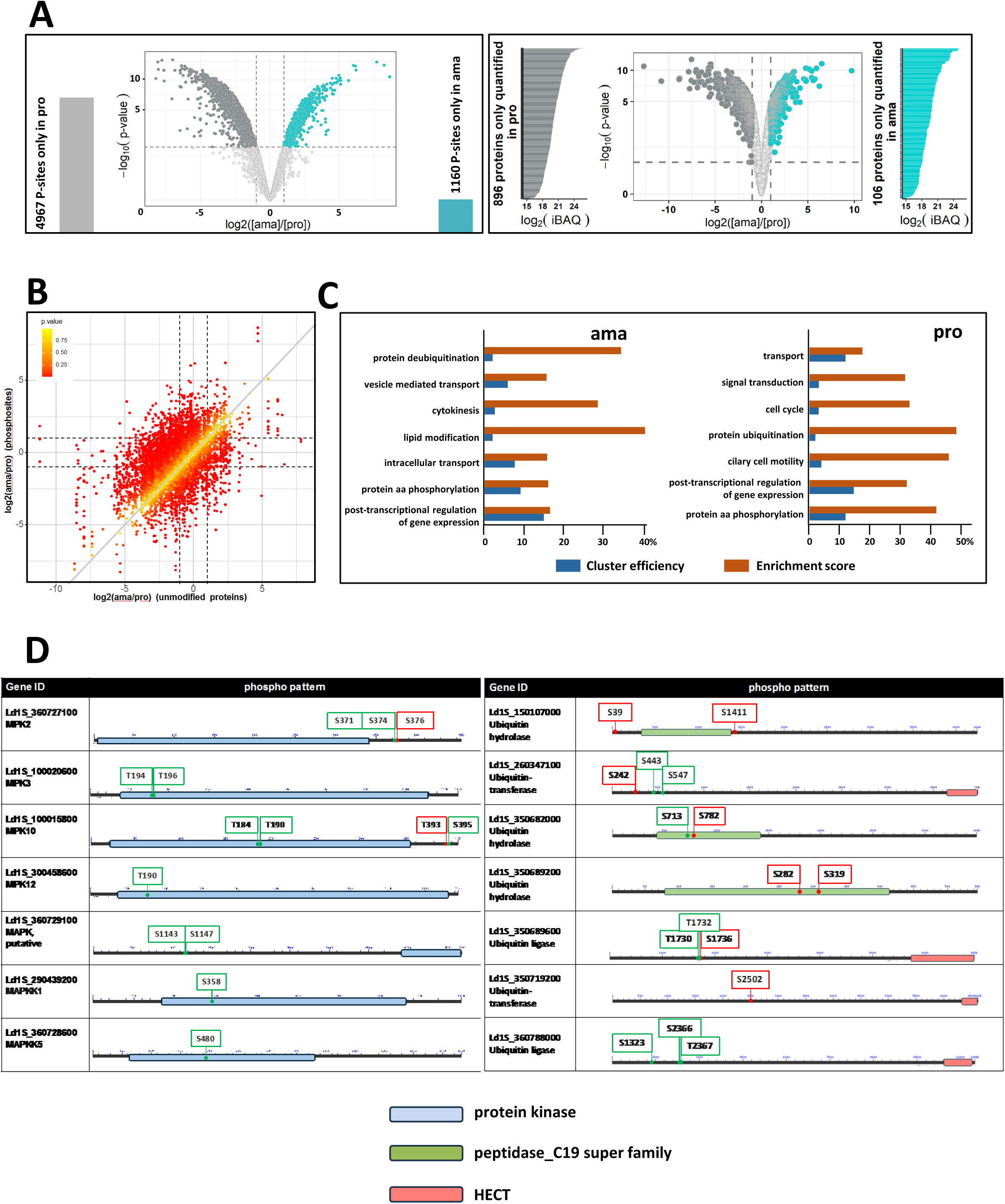
Stage-specific phospho-proteomic profiling. (A) Volcano plots corresponding to the total proteome (right panel) and phosphoproteome (left panel) analyses. Proteins and phosphosites (P-sites) only identified in one stage are presented at each side of the volcano plots. Cyan dots (ama) and grey dots (pro) represent signals with FC ≥ 2 and FDR < 1%. Light grey dots represent non-significant changes. (B) Relative phosphorylation change normalized to protein abundance. Ratio plot comparing the log2 fold changes in total protein abundance between ama and pro (x-axis) and phosphosite abundance in ama versus pro (y-axis). Dashed lines indicate a FC = 2. The color intensity of each dot reflects the p-value calculated for the relative phosphorylation change normalized to protein abundance as indicated in the graph. Confidence values were derived as described in Supplementary Methods. (C) GO term enrichment analysis for the category ‘biological process. Only stage-specific phosphosites were considered (i.e. sites that showed a significant increase in relative phosphorylation normalized to protein abundance, and sites that were only detected in one or the other stage, whether the protein was identified or not in the total proteome analysis). The histogram plots show ‘cluster efficiency’ in blue and ‘enrichment score’ in orange for the GO term enrichment analysis performed with the ama (middle panel) and pro (right panel) datasets. The main GO terms identified using the BiNGO plug-in in Cytoscape and associated with a BH p-value < 0.05 are shown. (D) Phosphorylation pattern of MAPK and MAPKK proteins (left panel) or Ubiquitin hydrolases and transferases (right panel) identified in the relative phosphorylation change normalized to protein abundance. The positions of the phosphorylated amino acid specific to ama (red) and pro (green) are indicated. Only phosphopeptides identified in the relative phosphorylation change normalized to phosphorylation level (see Table 11) and detected either in one stage or the other or quantified with a FC ≥ 2 were considered for the identification of the phosphosites.

We identified 877 phosphoproteins in amastigote samples carrying 2079 phosphosites of which 55.8% (1160) were uniquely identified at this stage, while 12.6% (565) showed a significant change in normalized phosphorylation levels compared to promastigotes (Figure S10, Table 11 sheets A to D). Conversely, 2054 phosphoproteins with 7095 phosphorylation sites were identified in promastigote samples, of which 70% (4967) were exclusively detected at this stage, while 1% (73) presented a significant change in normalized phosphorylation levels compared to amastigotes (Figure S10, Table 11 sheets F to I). These data reveal a surprising level of stage-specific phosphorylation in promastigotes, which may reflect their increased biosynthetic and proliferative activities compared to amastigotes, or may be a consequence of culture adaptation.

We next investigated the pathways regulated by phosphotransferase activities analyzing the phosphoproteomes of amastigote and promastigote for enriched biological processes (Figure 5C) (Table 11, sheets E and J). Stage-specific enrichment was observed in the pro dataset for ‘ciliary cell motility’, indicating that flagellar biogenesis and activity are not only regulated by increased protein abundance, but also by increased phosphorylation of flagellar proteins. Surprisingly, in both the ama and pro datasets, similar functions were enriched, including cell division (e.g. ‘cytokinesis’, ‘cell cycle’), signaling (e.g. ‘protein aa phosphorylation’, ‘signal transduction’), or post-transcriptional regulation of gene expression. Thus, aside stage-specific protein turnover (see above, Figure 4), differential phosphorylation represents yet another layer of regulation that contributes to the establishment of biological networks in *Leishmania* that share similar function in both ama and pro, but whose components are stage-specifically regulated at post-translational levels.

Analyzing stage-enriched, functional networks in further detail using the STRING applications of the Cytoscape software package revealed a series of potential regulatory interactions (Figure S11, Table 12): A first group of protein kinase substrates is represented by protein kinases themselves with more than 30 and 100 members of this large protein family identified in the ama and pro phosphoproteomes, respectively (Table 11, sheets D and H, see Figure 5D for phosphorylation pattern of some members of the MAP kinase pathway). An additional group of protein kinase substrates in both amastigote and promastigote phosphoproteomes include stage-specific phosphatases (18 in ama and 55 in pro) that can counteract protein kinase activities and even deactivate signaling cascades by dephosphorylation of protein kinase substrates (Figure S11, Table 11, sheets D and H). Furthermore, the stage-specific phosphoproteomes showed enrichment in functional networks linked to ribosomal biogenesis and RNA processing suggesting that protein kinases may control the establishment of stage-specific ribosomes. Finally, the identification of networks linked to proteasomal protein degradation in both ama and pro datasets not only further sustains the role of protein turnover in *Leishmania* stage differentiation, but uncovers a possible feedback loop between proteasomal activities controlling stage-specific degradation of protein kinases (see Figure 4), which in turn may control stage-specific ubiquitin ligases and deubiquitinating enzymes via differential phosphorylation (see Figure 5D for phosphorylation pattern of selected members of the Ubiquitin proteasomal system).

## Discussion

By applying a 5-layer systems analysis to spleen-derived bona fide amastigotes and culture-derived promastigotes, we identified a series of functionally related gene clusters that are co-regulated during *Leishmania donovani* stage differentiation. This co-regulation occurs at multiple levels, including transcript stability, proteasomal protein turnover, and protein phosphorylation. Our study underscores the complex molecular architecture of the *Leishmania* developmental process and provides novel insights into possible regulatory feedback loops that may coordinate stage-specific transitions. These results raise important questions regarding the integration of these regulatory networks during differentiation, and their contributions to environmental sensing and adaptive evolution. In the following we discuss these putative networks in the context of the current literature and propose experimental strategies for their downstream, mechanistic analyses.

A first series of possible regulatory networks proposed by our results could rely on recursive or self-controlling interactions, where components of a given pathway are regulated by the pathway itself (Figure 6A). In promastigotes for example, a co-regulated, functional gene cluster showing stage-specific increase in transcript abundance was defined by the GO term ‘post-transcriptional regulation of gene expression’. In the absence of stage-specific gene dosage changes (Figure 1) and the lack of transcriptional regulation of gene expression in *Leishmania* (104) the stage-specific expression changes of these transcripts thus is likely regulated itself at the post-transcriptional level. Such auto-regulatory feedback is common for RNA-binding proteins (RBPs), which play a central role in controlling mRNA stability, often targeting their own transcript (105). This helps keep protein levels stable (negative feedback) or create on/off switches for developmental processes (positive feedback) (106). In *T. brucei* <u>for example</u>, the stage-specific expression of RBPs, such as ZC3H22 or RBP9, is regulated at post-transcriptional levels (107), while other RBPs, such as TbZFP3 or RBP10, can control developmental transitions by stabilizing cohorts of mRNAs, thus defining developmental regulons (108, 109). In *Leishmania*, RBPs are likely to participate in comparable regulatory networks. These can be investigated through methods such as crosslinking and immunoprecipitation (CLIP) to map RBP binding to 3’ untranslated regions (3’UTRs), computational approaches to identify conserved regulatory sequence motifs, and loss-of-function analyses targeting either trans-acting factors (RBPs and their interaction partners) or cis-acting elements (e.g., RBP binding sites within 3’UTRs).

**Figure 6:**
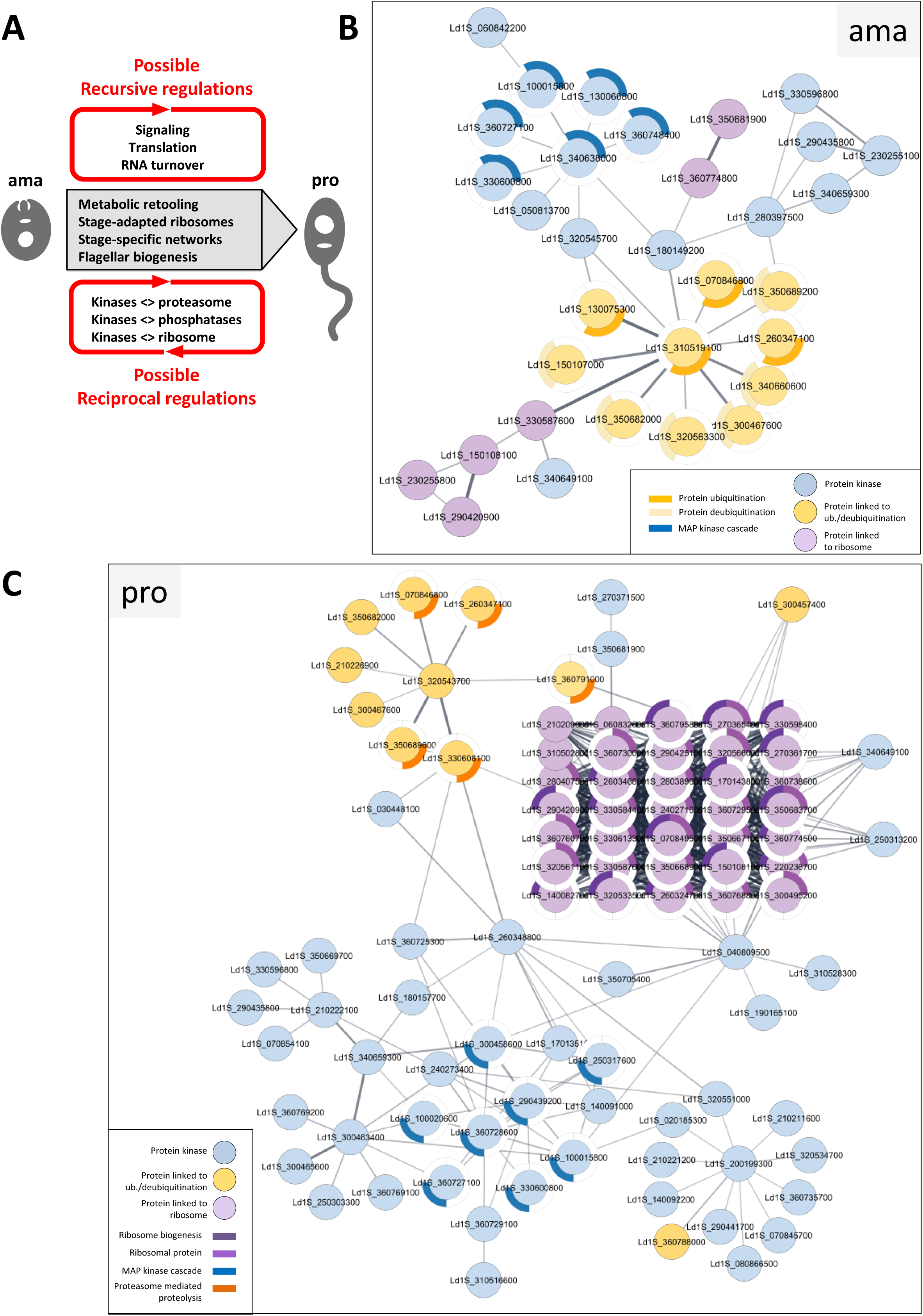
Network analyses. (A) Model of *Leishmania* gene expression regulation during stage differentiation. (B-C) Network analysis. Networks, restricted to the protein kinases (blue nodes), proteins implicated in ribosome biogenesis and ribosomal proteins (violet nodes) and proteins linked to ubiquitination/deubiquitination (orange nodes) identified based on normalized phosphorylation levels conducted in ama (B) or pro (C), were generated with the STRING plug-in of the Cytoscape software package using *L. infantum* orthologs and full STRING network with a confidence score cutoff of 0.4. Each node represents a phosphoprotein with the respective gene identifier indicated. GO terms associated to the proteins are represented by the colored segments around the nodes according to the legend shown in the graph. Only GO terms associated with a p-value < 0.05 were considered.

Our data propose a similar recursive feedback loop that seems to act on the level of protein translation, which was revealed by a paradox: while amastigotes showed increased abundance for 38 transcripts annotated for the GO term ‘ribosome biogenesis’ (Table 4, sheet E), the corresponding proteins showed reduced abundance in amastigotes compared with promastigotes (Figure 3C). This dissociation between RNA and protein abundances indicates that ribosomal components themselves may be regulated at translational levels. Increased ribosomal protein abundance in promastigotes could satisfy the requirement of increased translation capacity at this fast-growing stage (31, 110). In contrast, increased mRNA abundance of ribosomal components in amastigotes may indicate stage-specific mechanisms of mRNA storage (111) that may allow to jump start promastigote differentiation and accelerated growth following exposure to low temperature and neutral pH. This possibility is supported by our proteomics investigations of the early differentiation process, revealing ribosomal biogenesis as one of the first pathways induced over the initial 18h during the amastigote-to-promastigote developmental transition (Figure S6).

Differentiation to promastigotes was associated not only with quantitative changes of ribosomal components as indicated by co-regulated expression of ribosomal proteins and translation factors (see Table 8), but also seems to affect ribosome structure itself as indicated by stage-specific expression changes of gene sets enriched in the GO terms ‘RNA methylation’ (8 methyltransferases, including the rRNA methyltransferases Ld1S_350672300 and Ld1S_350694000) and ‘pseudouridine synthesis’ (6 pseudouridine synthases and the H/ACA snoRNP Nop10 (Ld1S_360722900) (Table 8, sheets D to H). The differential expression of snoRNAs (see Figure 2C and Table 5) and the developmentally regulated changes in rRNA modification that can affect ribosome activity (34) (Figure 2D) indeed support the existence of such stage-specific ribosomes. These results are in line with our recent reports providing indication for specialized ribosomes in *Leishmania* adaptation, in which we correlated changes in rRNA pseudouridylation to *L. donovani* fitness gain in culture (31, 66). Cryo-EM analysis of purified ribosomes from amastigotes and derived promastigotes, combined with targeted deletion of individual snoRNA genes, will be critical to uncover stage-specific structural and functional adaptations of such stage-specific ribosomes.

Finally, a third possible recursive feedback loop seems to act at the post-translational level involving members of various protein kinase (PK) families, which not only showed stage-specific expression, but also represented a main class of phosphorylation substrates themselves. This interaction may reflect stage-specific signaling cascades, where downstream kinases are activated through phosphorylation by upstream kinases, such as exemplified by the MAP kinase pathway (103, 112). Indeed, 11 members of the MAP kinase family showed stage-specific changes in their phosphorylation pattern (Figure 5D), 9 of which were only detected in the promastigote phospho-proteome and were previously associated with this stage, being implicated in flagellar biogenesis (MPK9 and MPK3) or metacyclogenesis (MPK4) (98, 102, 113). In contrast, the two MAP kinases MPK2 and MPK10 - previously linked to amastigote differentiation and virulence (99, 102) - showed an amastigote-specific increase in protein abundance and relative phosphorylation change for defined phosphorylation sites. In *L. mexicana*, such a recursive PK feedback loop has been identified between the interacting protein kinases LmxMKK and LmxMPK3, which cross-phosphorylate each other to regulate flagellar length (113). To further dissociate the complex regulatory relationships between parasite PKs, protein-protein interaction maps should be established (e.g. co-IP, proximity labelling), combined with in vitro kinase assays to reveal direct kinase-substrate relationships.

Most PKs have pleiotropic effects across biological systems, due to their ability to phosphorylate a wide range of functionally unrelated substrates (114). Our analysis of relative phosphorylation changes normalized to protein abundance confirms such pleotropic function in *Leishmania* and suggests parasite PKs at the center of a second form of possible feedback loop defined by reciprocal interactions, where components of two or more regulatory pathways may cross-control each other (Figure 6A). For example, as judged by changes in phosphorylation abundance, it seems that stage-specific PK activities act on a series of phosphatases, which in return may modify kinase activities by dephosphorylation (Table 11, sheets D and G). Such reciprocal regulation between both enzyme families is well-established across many eukaryotic systems, for example between mammalian ERK1/2 and the MAPK phosphatase DUSP6/MKP-3 (115), or yeast CDK1 and the phosphatase Cdc25, controlling mitotic entry (116). In addition, 47 differentially phosphorylated proteins were associated with ‘Translation’, some of which have been linked in *Leishmania* to translational control, cell cycle regulation or infectivity (29, 117–121). This opens the possibility of yet another reciprocal feedback loop, where phosphorylation of ribosomal components could fine tune translational control, which in turn may affect the expression of protein kinases themselves (122–130).

Finally, our results suggest the possibility of a third reciprocal feedback loop between protein kinases and proteins of the proteasomal system that may govern *Leishmania* stage transitions. Previous genetic analyses demonstrated an essential role for various components of this system in promastigote-to-amastigote transition and intracellular survival, including *L. mexicana* ubiquitin conjugating (E2) enzymes (UBC1/CDC34, UBC2 and UEV1), the E3 ubiquitin ligase HECT2 or various deubiquitinases (e.g. DUBs 4, 7, 13) (91, 92). By applying the irreversible proteasome inhibitor lactacystin, we extended this role to the reverse process - amastigote-to-promastigote differentiation - demonstrating that proteasomal activities are critical for both directions of the developmental cycle (Figure 4). The constitutive expression of many proteasomal components across both amastigote and promastigote stages supports a role of phosphorylation in regulating stage-specific proteasomal activities, considering that phosphorylation of DUBs has been linked to changes in their stability, localization, specificity or catabolic activity (131, 132). We indeed identified 11 DUBs and 10 ubiquitin ligases/transferases showing stage-specific phosphorylation at specific residues (Table 11, sheets D and I). This correlated with the stage-specific proteasomal degradation of 14 PKs (Figure 4F, Table 9, sheet G), including two previously implicated in *Leishmania in vivo* fitness (Repressor of differentiation kinase 1, Ld1S_320540600; differentiation inhibitory kinase, Ld1S_280397500) (102). These findings support the existence of a possible kinase/proteasome reciprocal feedback loop in early parasite development. The experimental validation of such regulatory interactions will require in-depth biochemical, molecular, and systems biology approaches involving global kinase-substrate mapping (114), in vitro reconstitution assays using recombinant proteins to demonstrate reciprocal kinase and proteolytic activities, and genetic perturbation test to reveal the biological impact of the regulatory interaction.

The complexity of regulatory interactions between protein kinases and the proteasomal pathway is further increased by observations showing that substrate phosphorylation can mediate subsequent ubiquitination and degradation, while ubiquitination of MAP kinase family members can regulate kinase activity and localization rather than promote degradation (139–141). Future experiments simultaneously analyzing changes in both phosphoproteome and ubiquitome during *Leishmania* stage-differentiation and making precision mutants that lack the ubiquitination or phosphorylation sites may help elucidate this complex regulatory interaction and identify individual proteins and their PTMs that drive parasite development

In conclusion, our data represent an important novel resource to experimentally assess the regulatory landscape of *Leishmania* stage development. We propose genetic feedback control as a central mechanism in parasite differentiation, with stage-specific interactions between genes and their products likely adapting the parasite phenotype to the environmental changes encountered inside its vertebrate and invertebrate hosts. In contrast to the major phenotypic shift observed between mammalian-stage amastigotes and insect-stage promastigotes, each developmental stage is stably maintained within its respective host even in the wake of environmental fluctuations. Such phenotypic robustness is known to rely on highly inter-connected, stage-specific regulatory networks (142), which in the case of *Leishmania* could implicate recursive and reciprocal genetic interactions between protein translation, protein degradation and protein phosphorylation (Figure 6B and C). Our results define *Leishmania* differentiation as an excellent model system to study how alternations between phenotypic plasticity (i.e. developmental transitions) and robustness (stable maintenance of stages) shape microbial fitness, and provides a powerful experimental framework for future studies on the emergence of stable, adaptive biological traits through feedback regulation.

## Acknowledgments

We thank the CEA-CNRGH for its contribution to the sequencing costs and all the CEA-CNRGH staff who performed sample preparation and sequencing for their excellent technical assistance. We acknowledge the help of the HPC Core Facility of the Institut Pasteur for this work. We thank Georges Haustant and Caroline Proux from the Biomics Platform, C2RT, Institut Pasteur, Paris, France for the library preparation and the sequencing, Hugo Varet and Rachel Legendre from Institut Pasteur, Université Paris Cité, Biostatistics and Bioinformatics Hub, Paris, France for their contribution to the RNA seq data analysis. We thank Karim Sébastien from the animal facility of the Institut Pasteur for his commitment in the care of our animals.

This work was supported by the Agence Nationale pour la Recherche Labex ‘Integrative Biology of Emerging Infectious Diseases’ contract ANR-10-LABX-62-IBEID and Labex ‘French Alliance for Parasitology and Health Care’ contract ANR-11-LABX-0024 (GFS, PP), the France Génomique National infrastructure, funded as part of the « Investissements d’Avenir » program managed by the Agence Nationale pour la Recherche contract ANR-10-INBS-09 (GH, HV, RL, CP), and the ERC SYNERGY project DecoLeishRN, Grant agreement ID: 101071613.

## Sup figure legends

**Figure S1:**
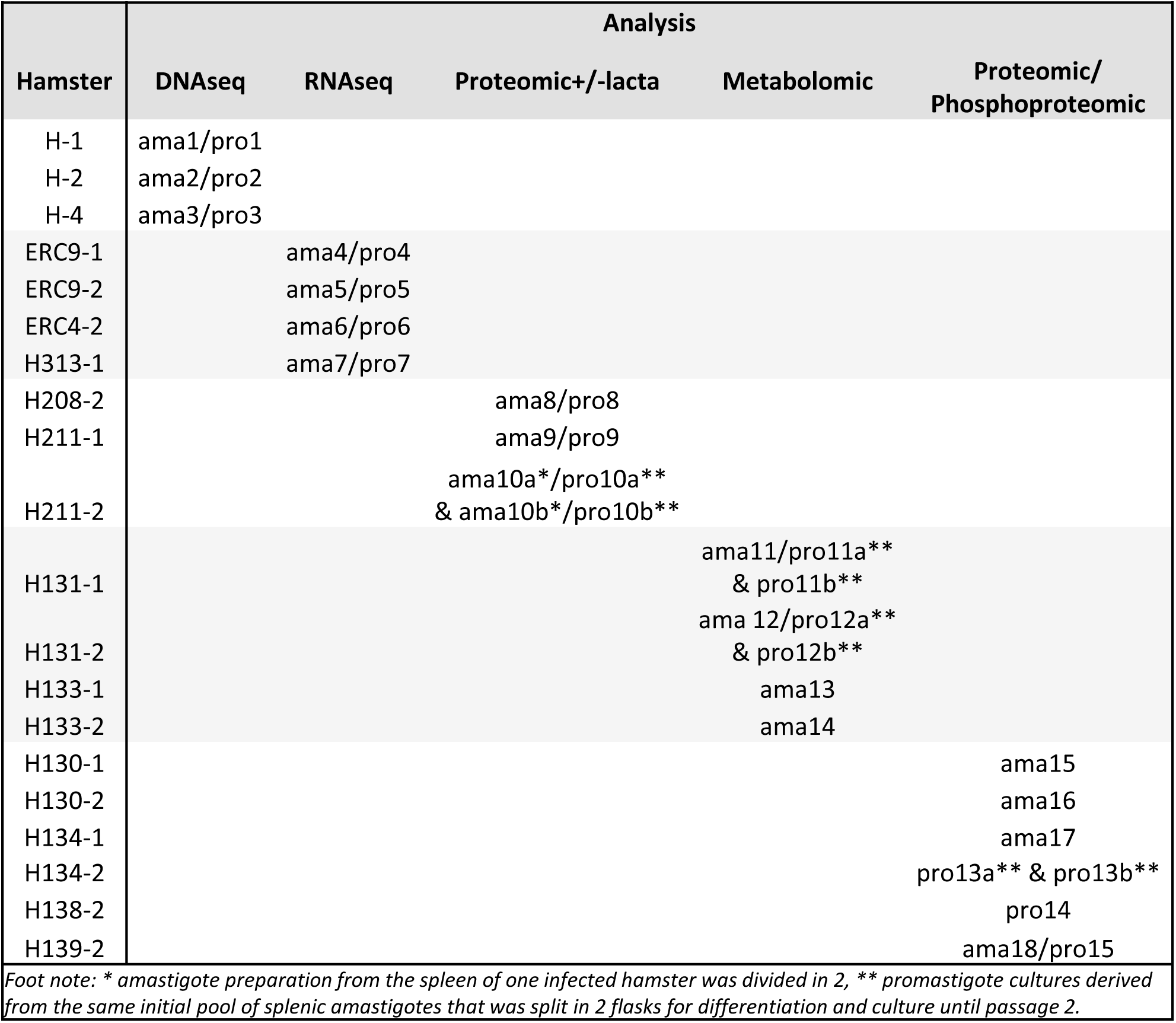
Overview of the samples used in this study. Each *L. donovani* infected hamster is identified by the cage number, as are the hamster-derived amastigotes (ama) and corresponding promastigotes (pro).

**Figure S2:**
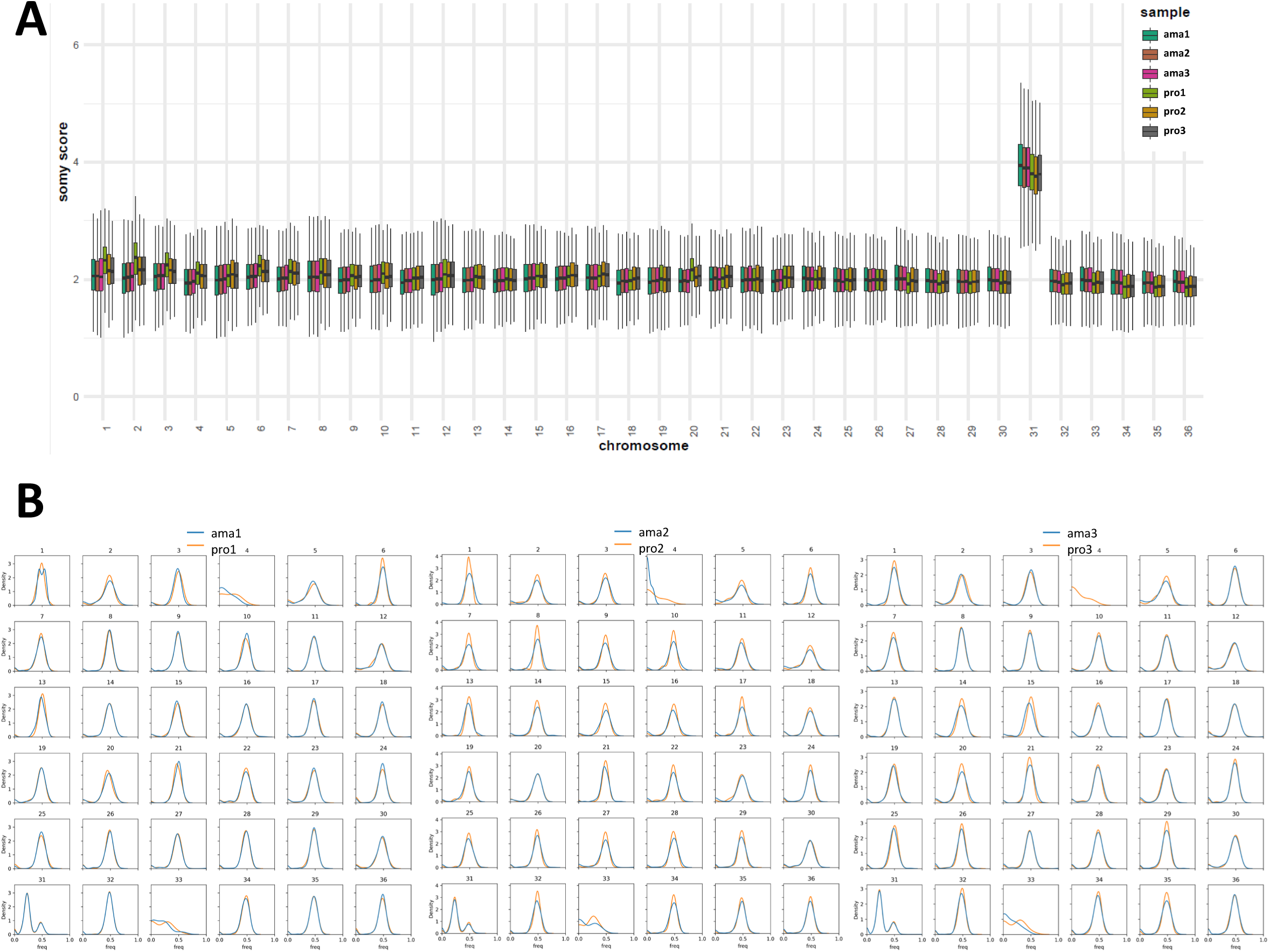
(A) Box plots representing the median somy score for all the chromosomes. Each box corresponds to one of three biological replicates for ama and pro as indicated in the graph. Sample identifiers are detailed in the legend of Figure 1A and Figure S1. (B) Frequency distribution plots representing, for each biological replicate and each chromosome, a kernel density estimate of the distribution of SNP frequencies in the amastigote and promastigote sample.

**Figure S3:**
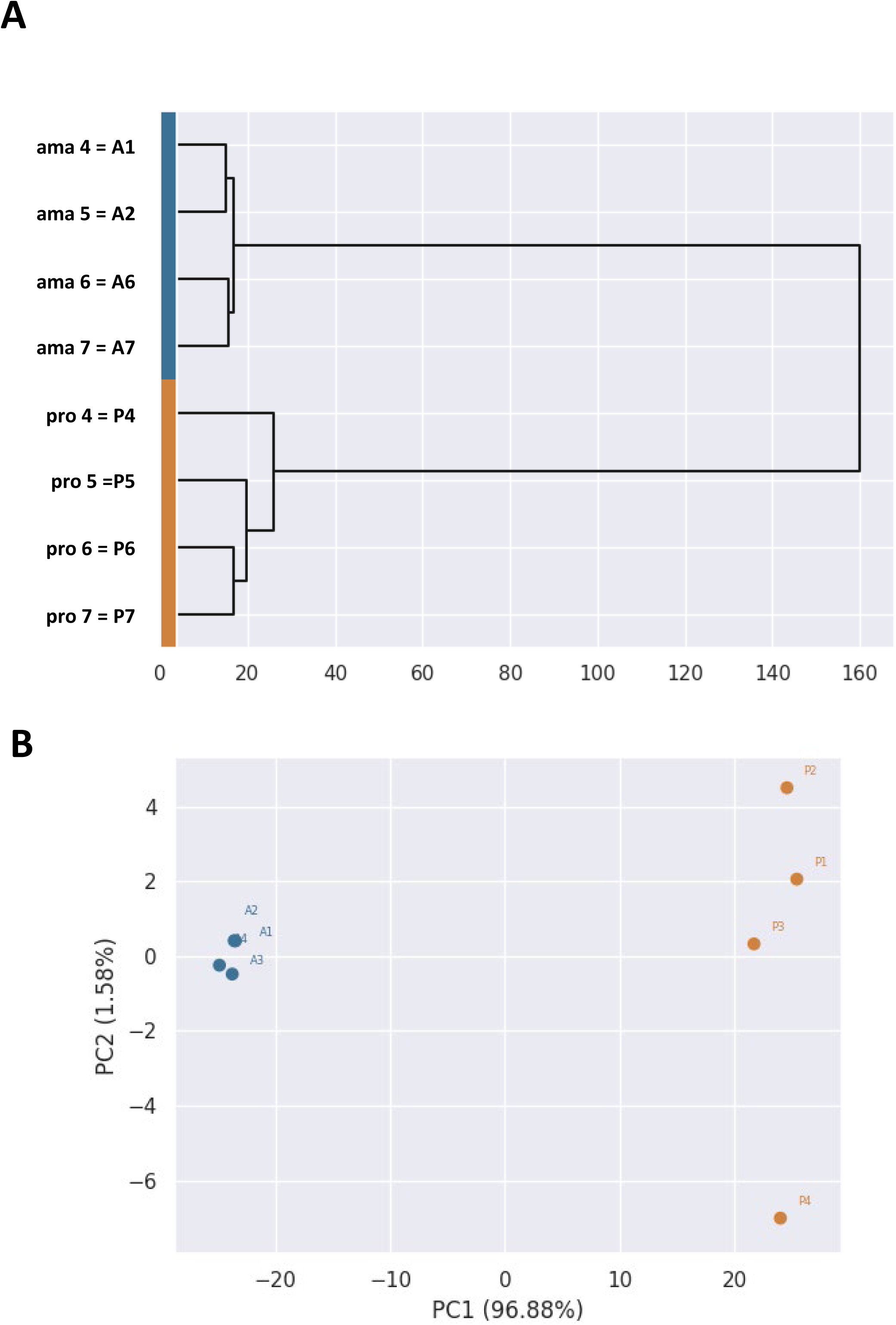
(A) Hierarchical clustering of the samples according to RNA-seq expression profiles. The dendogram is built using the Ward method. The normed data was log-transformed first and at max 5,000 most variable features were selected. (B) Principal component analysis. The two main components are represented. Ama and pro samples correspond to blue and orange dots respectively.

**Figure S4:**
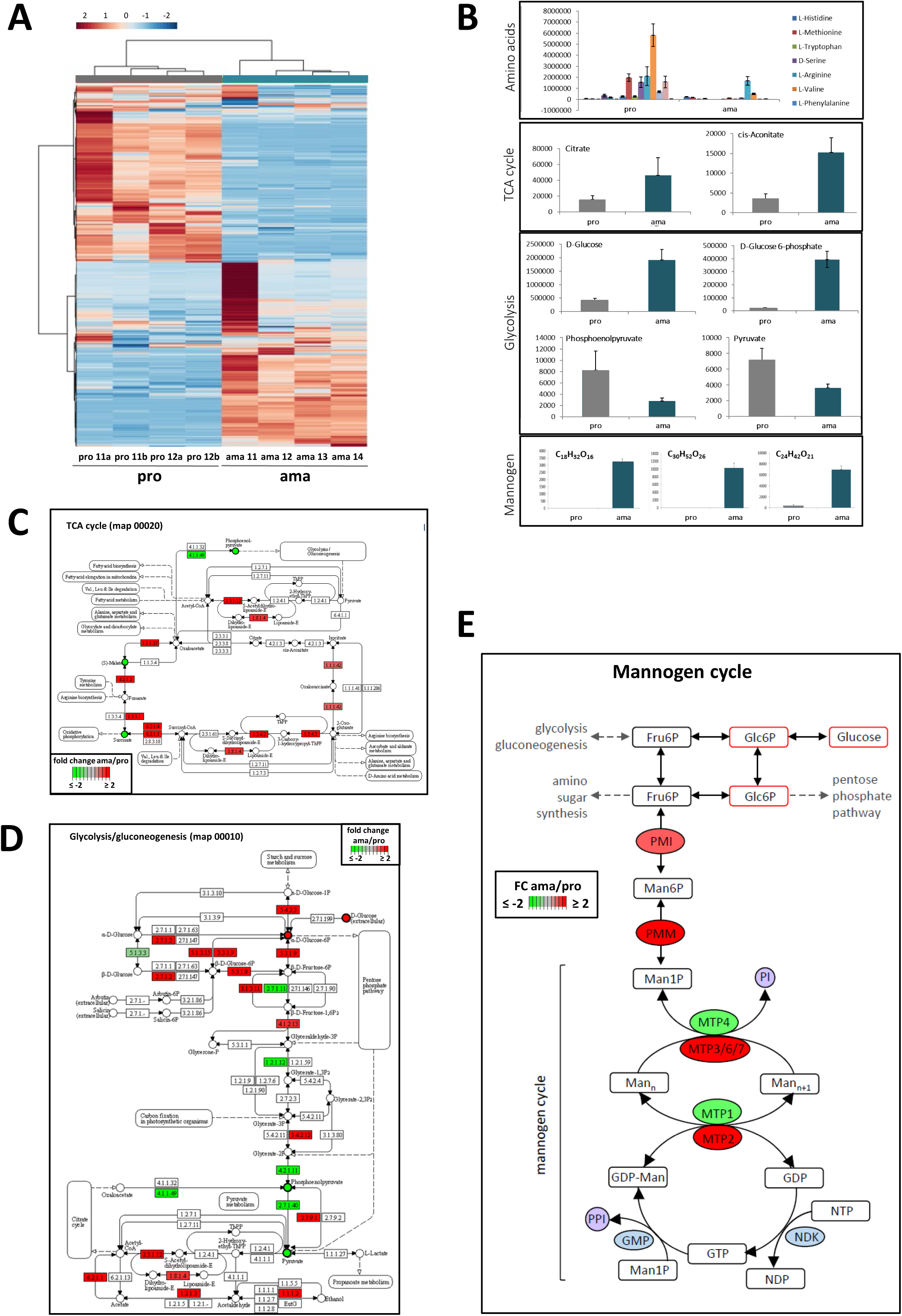
(A) Cluster analysis of metabolites showing differential abundance in ama and pro samples. (B) Quantification of amino-acids or metabolites involved in glycolysis, TCA cycle and mannogen cycle. The y-axis represents the intensity for each metabolite as detected by MS. (C-D) Correlation between metabolomic and proteomic analyses for TCA cycle (C) and Glycolysis/Gluconeogenesis (D). Fold changes in proteins abundance (squares) and metabolite abundance (circles) with adjusted p-values of respectively < 0.01 and < 0.05 were projected on the respective KEGG maps. The color intensity reflects the fold change value as indicated by the legends in the graphs. (E) Representation of the mannogen pathway. Metabolites are represented by a square, with red-lined squares corresponding to the metabolites that accumulate at the ama stage. Proteins are represented by a circle, with fold changes in protein abundance between ama and pro datasets indicated by color and intensity (red, increase; green, decrease).

**Figure S5:**
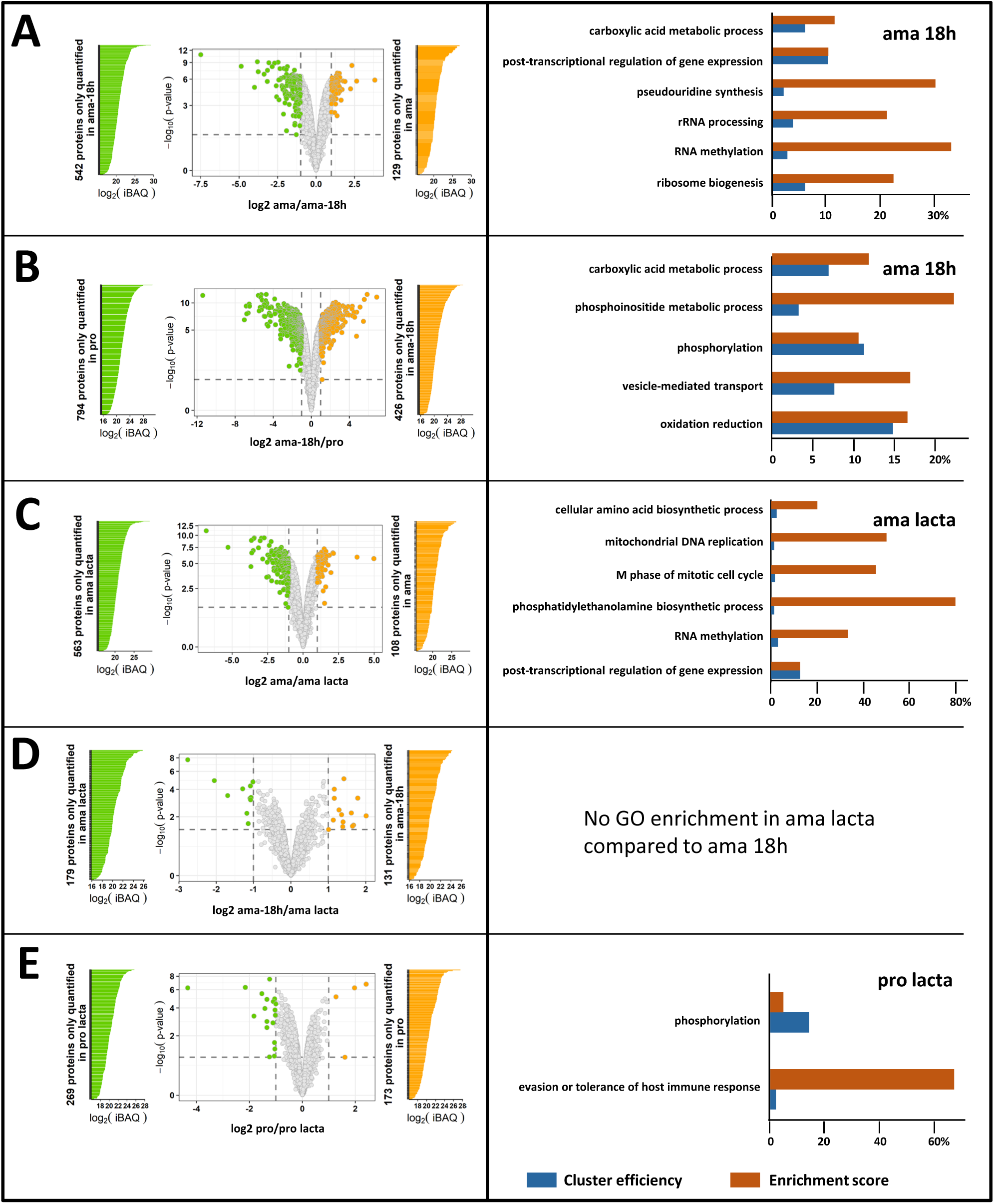
Label free quantitative proteomic analyses in presence or absence of lactacystin. (A) to (E) Volcano plots representing changes in protein abundance for the different comparisons (left panels) and GO term enrichment analyses for the indicated sample (right panels). Proteins identified by at least two peptides in at least three out of four biological replicates were considered. For volcano plots, colored dots indicate values with FDR < 0.01 and FC ≥ 2 (see Table 8). The light grey dots indicate non-significant expression changes. The colored bars indicate unique protein identifications in one (green) or the other (orange) condition with relative abundance indicated by the iBAQ value (left panels). Proteins quantified in only one condition or showing a significant increase in abundance (FC ≥ 2 and adjusted p-value < 0.01) were used to perform the GO analysis for the category ‘biological process’. The histogram plots show ‘cluster efficiency’ in blue and ‘enrichment score’ in orange (right panels) for a selection of GO terms identified using the BiNGO plug-in in Cytoscape and associated with a BH p-value < 0.05. (A) ama vs ama-18h; (B) ama-18h vs pro; (C) ama vs ama lacta; (D) ama-18h vs ama lacta; (E) pro vs pro lacta.

**Figure S6:**
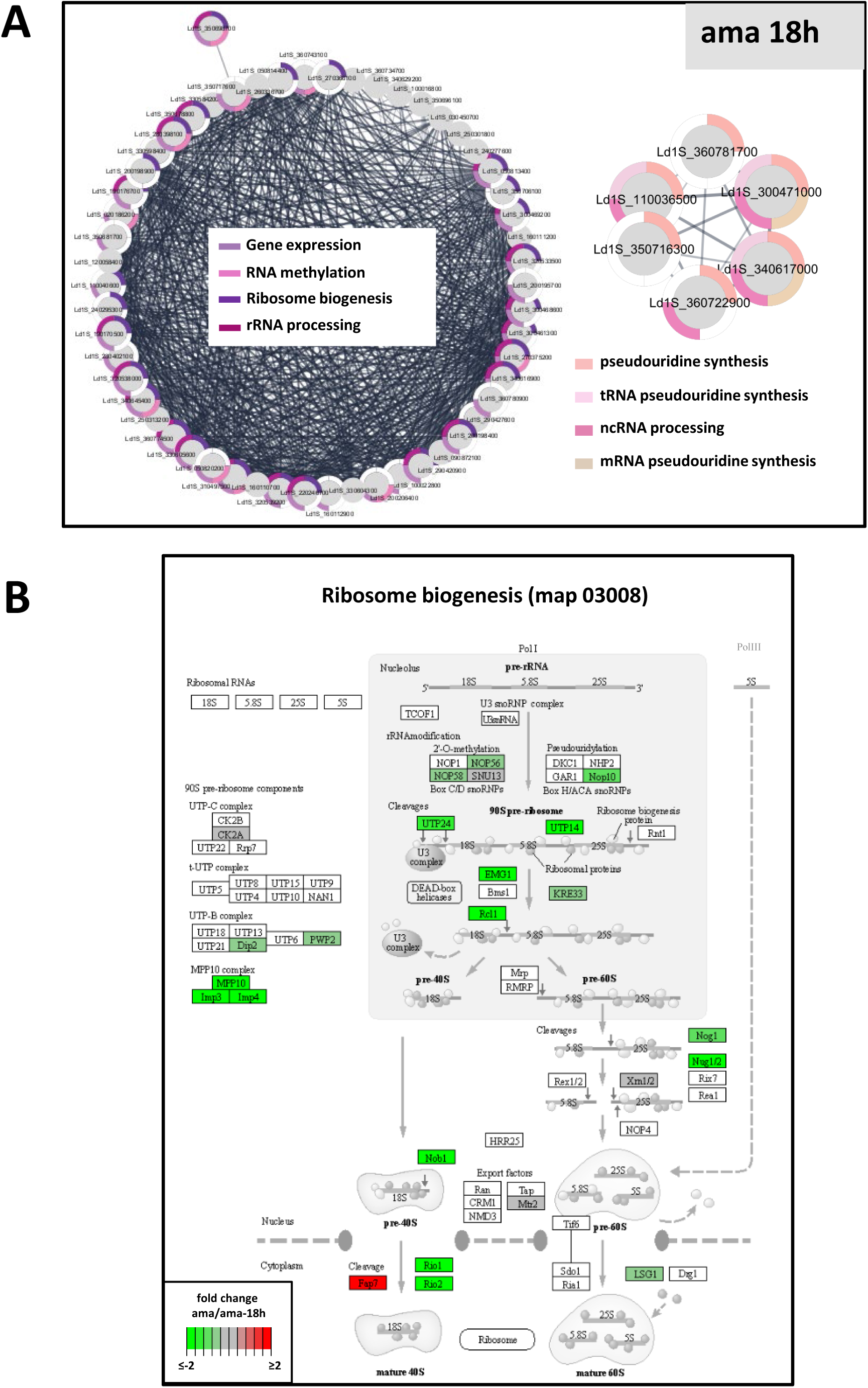
(A) Functional network enrichment analysis for proteins showing increased abundance in ama-18h compared to ama. The network was generated with the *L. infantum* orthologs using the STRING plug-in of the Cytoscape software package and considering the full STRING network with a confidence score cutoff of 0.4. The GO terms associated with each protein are represented by the colored segments according to the legend. Only GO terms associated with a p-value < 0.05 were considered. (B) KEGG map for ribosome biogenesis showing changes in abundance protein between ama and ama-18h. Only proteins with an adjusted p-value < 0.01 were considered. Differential abundance is indicated by the color and its intensity (red, increase; green, decrease).

**Figure S7:**
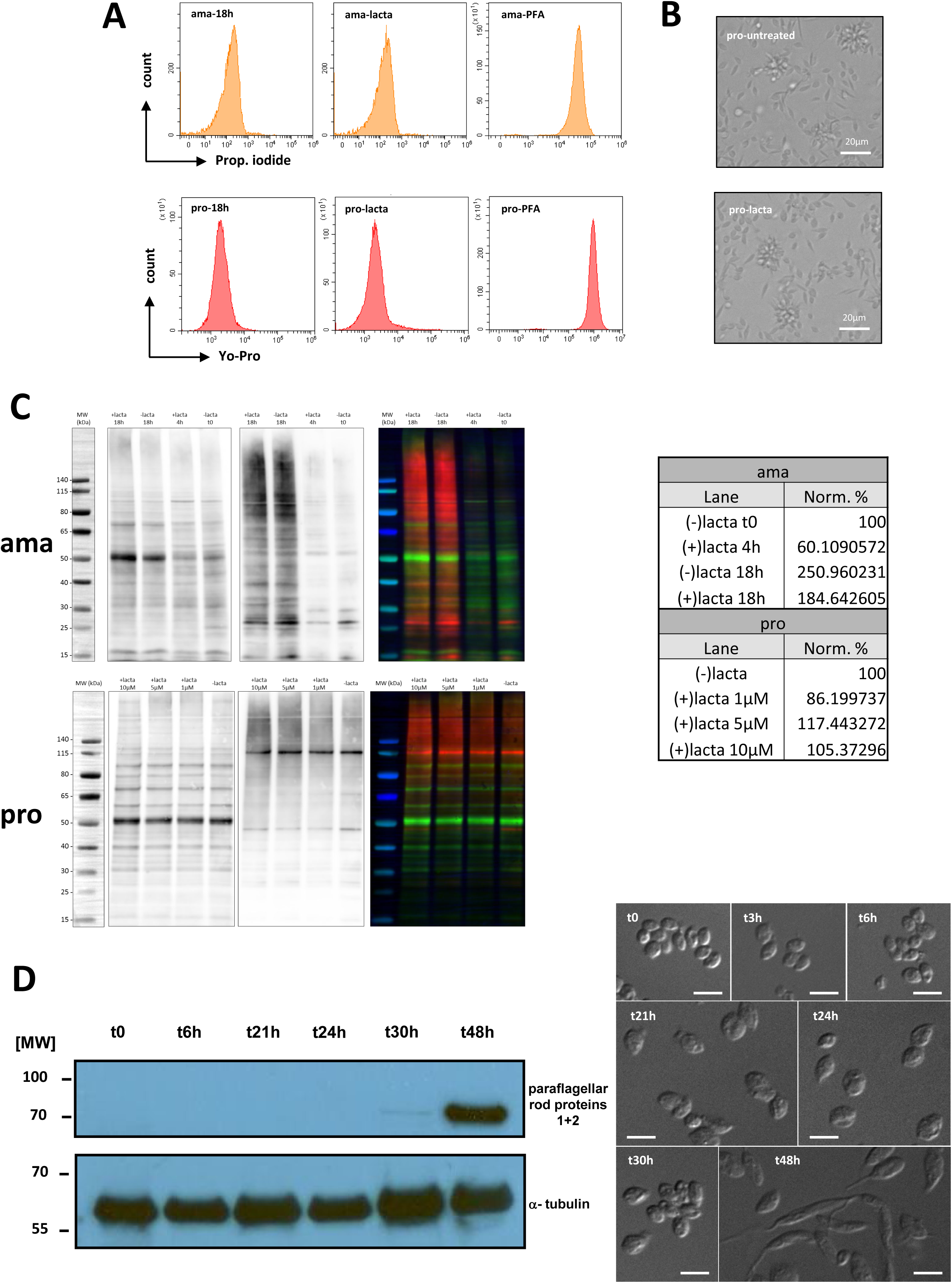
(A) Viability assessment of the indicated amastigote (upper panel) and promastigote (lower panel) samples after lactacystin treatment by FACS analysis using propidium iodide or YoPro staining. (B) Microscopic images of promastigotes at 18h in absence (upper image) or presence of 10 µM of lactacystin (lower image). (C) Western blot analysis of protein extracts obtained from ama (upper image) and pro (lower image) in presence or absence of lactacystin treatment. 10 µg of proteins were labelled with Cy5, separated on NuPAGE 4-12%, and transferred onto PVDF membrane. Membranes were subsequently incubated with a monoclonal antibody against ubiquitin and revealed with HRP-conjugated secondary antibodies. Images of Cy5-labelled proteins (left images), ubiquitinated proteins (middle images) and an overlay of both images (right images, Cy5-labelled proteins in green, ubiquitinated proteins in red) are presented. The table shows the relative quantification after normalization using the Cy5-labelled proteins of untreated samples as a loading control. (D) Western blot analysis of protein extracts obtained at the indicated time points after *in vitro* differentiation from hamster-derived amastigotes to promastigotes. 10 µg of protein were separated on NuPAGE 4-12%, transferred onto PVDF membrane subsequently incubated with antibodies against paraflagellar rod proteins 1 and 2 and alpha-tubulin, and revealed with HRP-conjugated secondary antibodies (left image). Micrographs showing hamster-derived amastigotes during *in vitro* differentiation to promastigotes (right image). Parasites were collected at the time points indicated, fixed in 4% PFA, and seeded on poly-L-lysine-treated coverslips. Parasites were observed with a 63x oil immersion objective and images were processed with AxioVision Rel.4.8 software. Differential interference contrast (DIC) microscopy was performed with Axioplan 2 imaging microscope using a 63x oil immersion objective, Axiovision software and AxioCam MRm camera (Carl Zeiss). The scale bars correspond to 1 µm.

**Figure S8:**
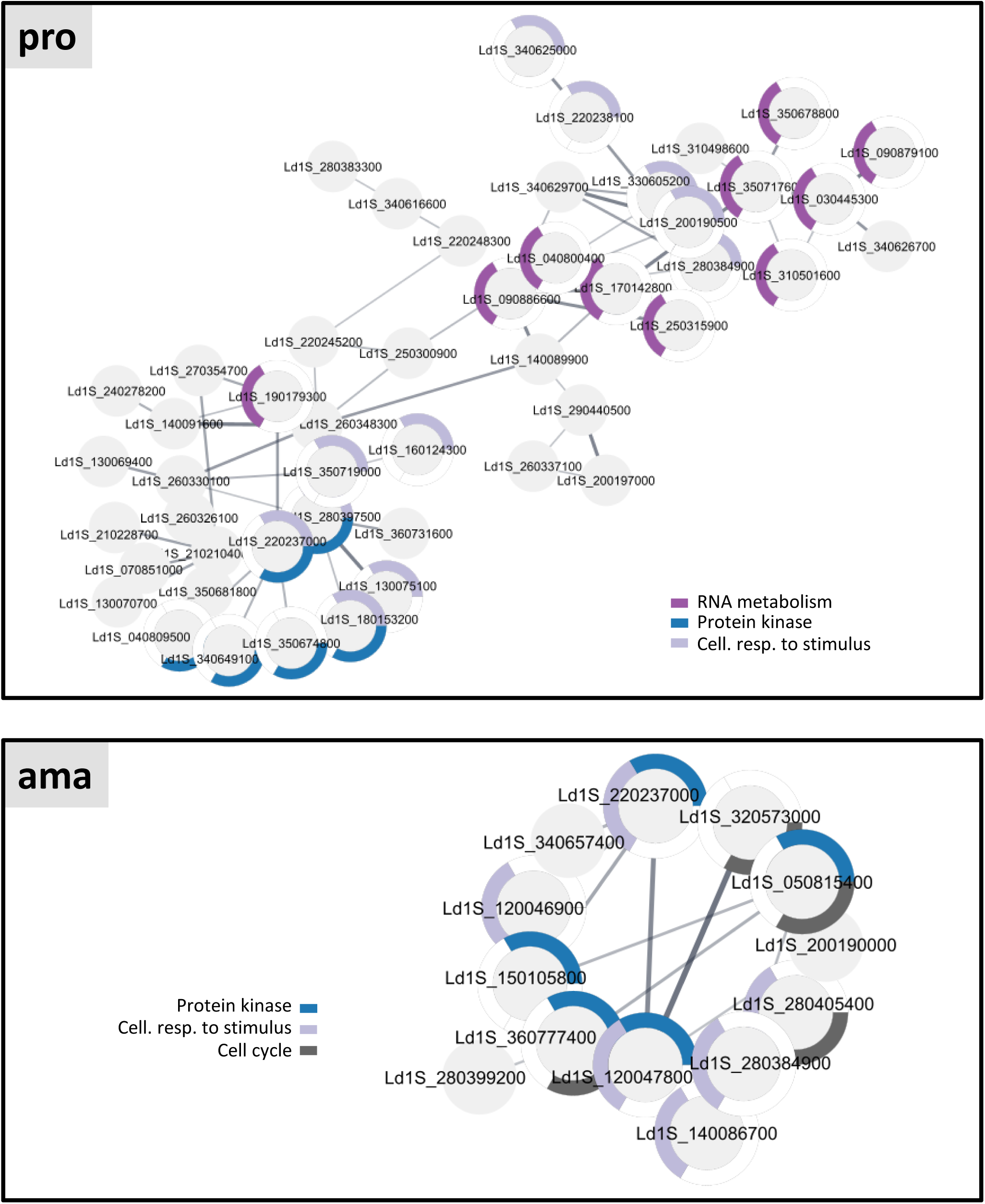
Functional network enrichment analysis for the proteins stabilized in presence of lactacystin in promastigote (pro) and amastigote (ama) datasets. The network was generated with the *L. infantum* orthologs using the STRING plug-in of the Cytoscape software package and considering the full STRING network with a confidence score cutoff of 0.4. The GO terms associated with each protein are represented by the coloured segments according to the legend. Only GO terms associated with a p-value < 0.05 were considered.

**Figure S9:**
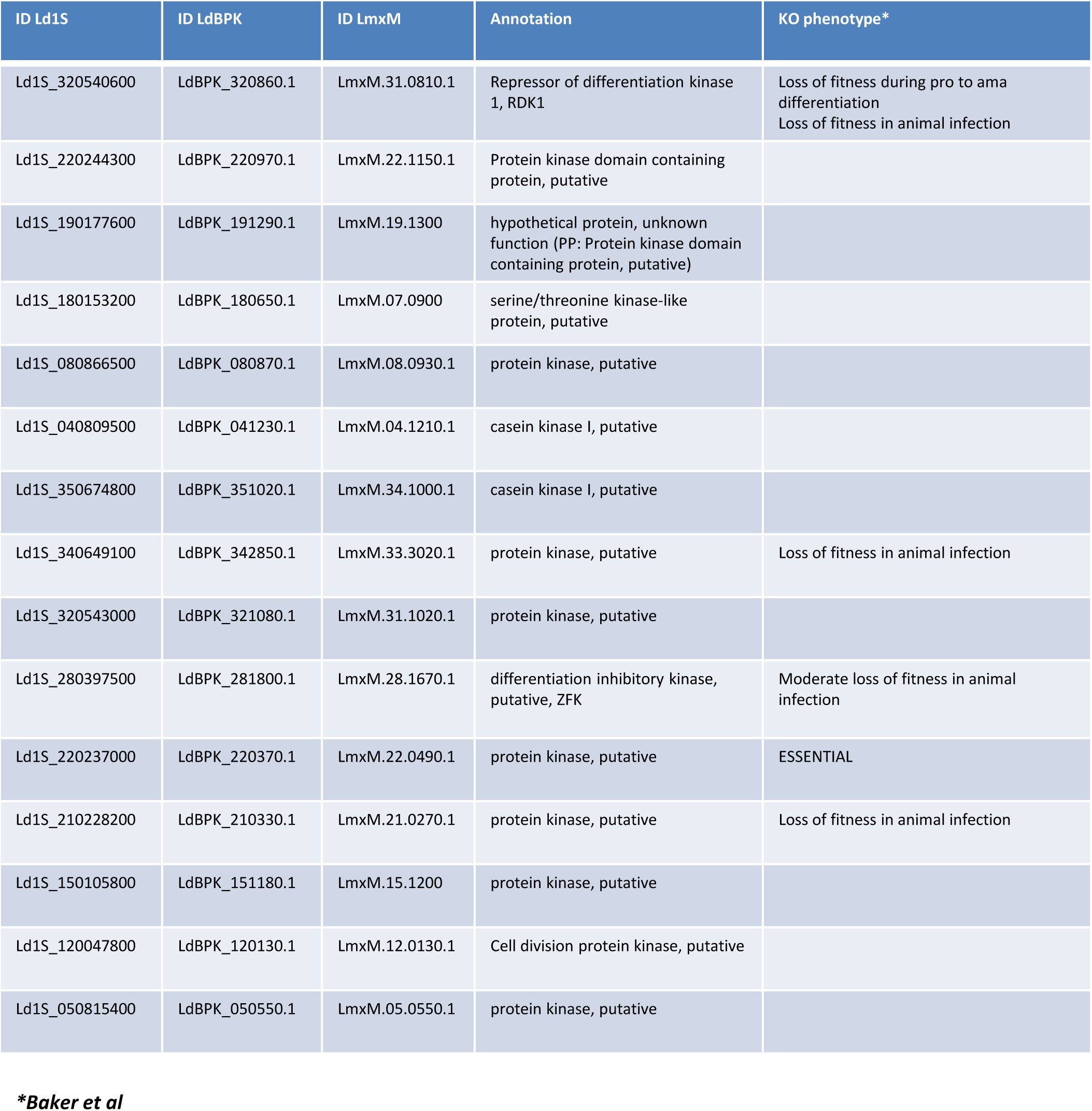
List of protein kinases stabilized in presence of lactacystin. Their corresponding orthologs in *L. donovani* strain LdBPK and *L. mexicana* are indicated, as well as the knock-out phenotype as published by Baker et al (2021) (102).

**Figure S10:**
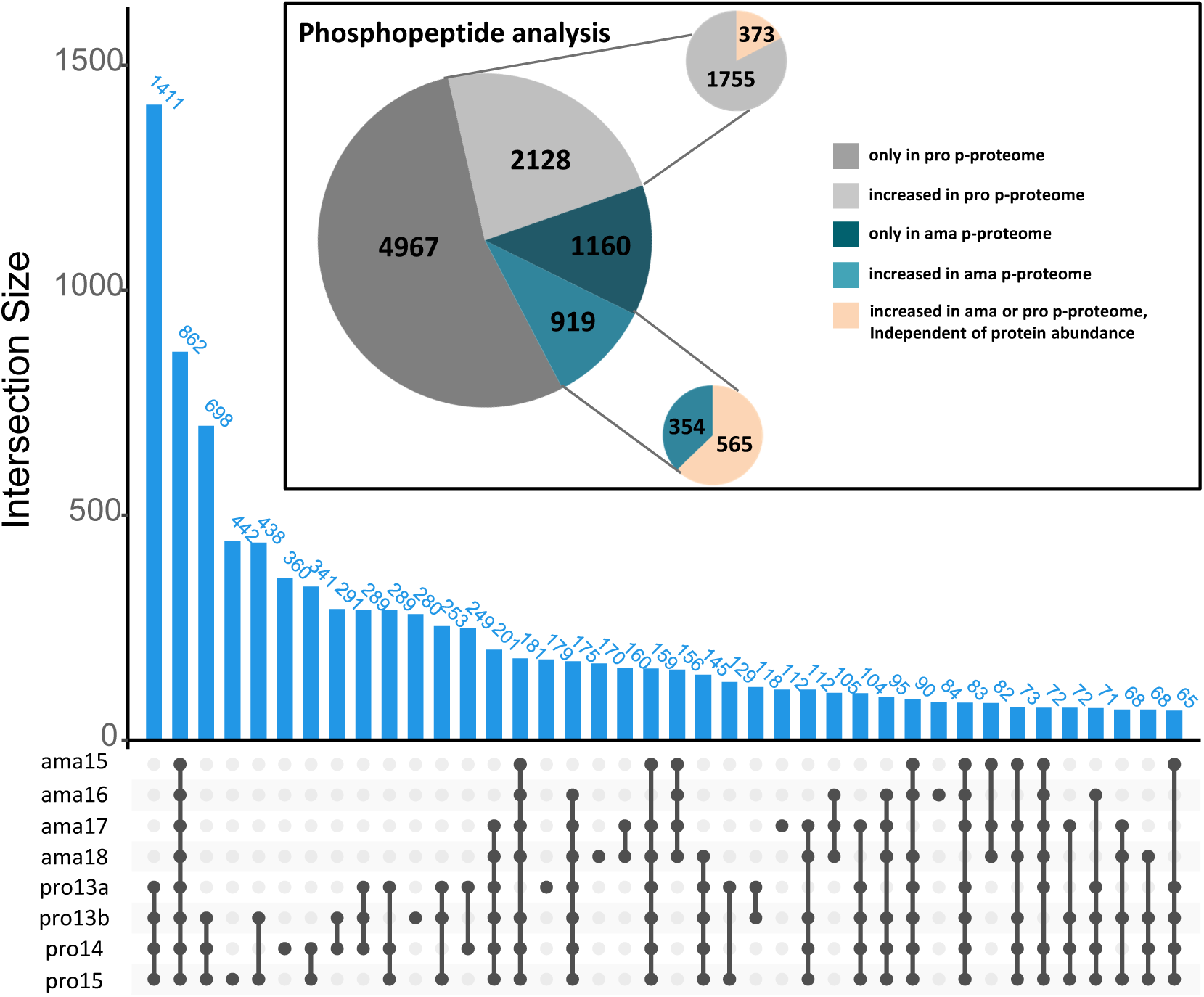
Upset plot representing the number of phosphopeptides (y-axis, blue histograms, numbers are indicated) and the samples in which they were quantified (x-axis). Pie chart of the phosphopeptide analysis showing the distribution of the phosphorylation sites according to the condition in which they were quantified as indicated in the legend of the graph.

**Figure S11:**
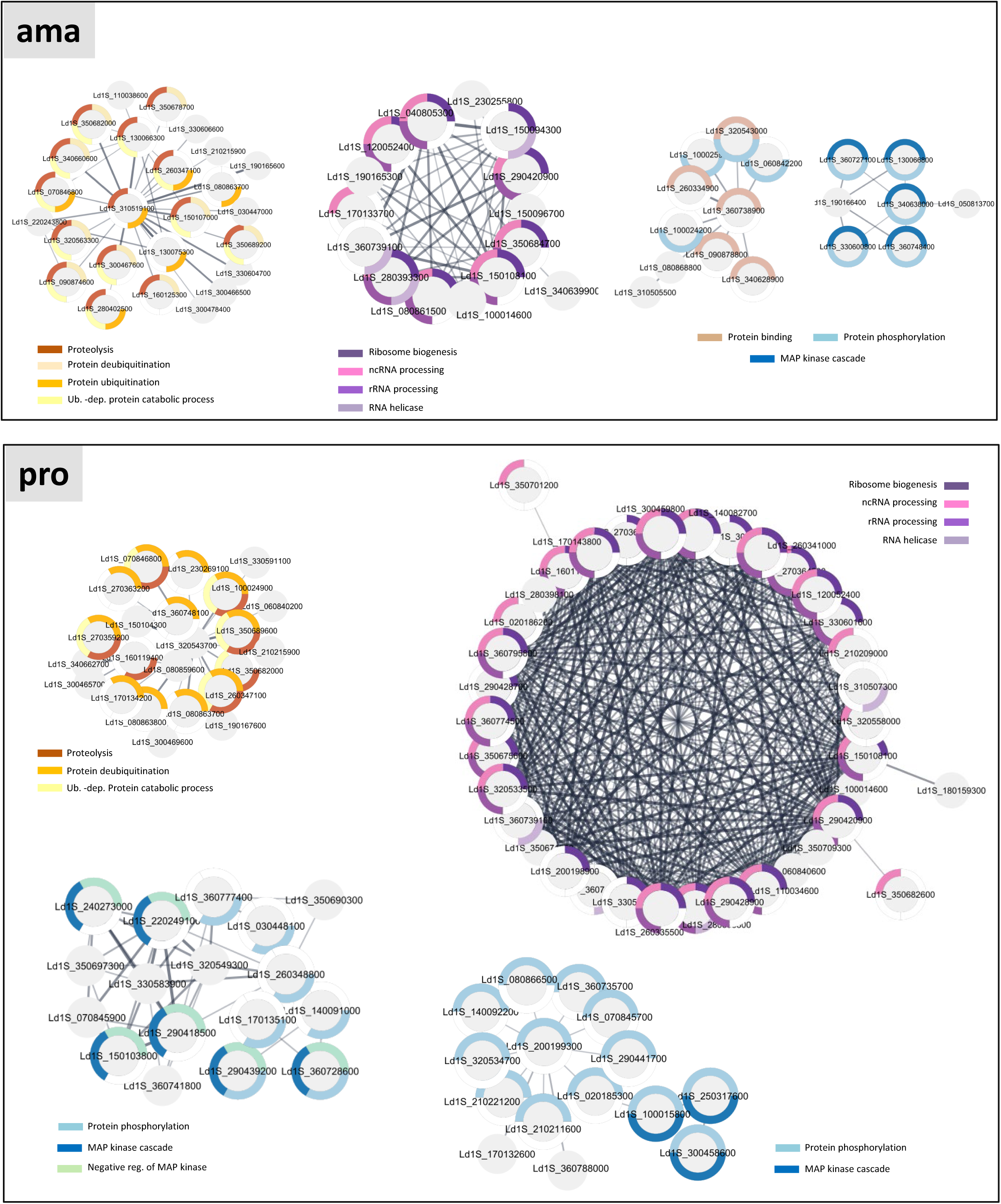
Functional Network analysis. Networks for proteins that show unique or increased phosphorylation in ama (upper panel) and in pro (lower panel) datasets were generated with the STRING plug-in of the Cytoscape software package using *L. infantum* orthologs and considering the full STRING network with a confidence score cutoff of 0.4. The nodes represent phosphoproteins associated with one or more GO terms (listed in the graph), with the respective gene identifier indicated. The GO terms associated to the proteins are represented by the colored segments around the nodes according to the legend shown in the graph. Only GO terms associated with a FDR < 0.05 were considered.

## REFERENCES

1. Hollin T, Le Roch KG. From Genes to Transcripts, a Tightly Regulated Journey in Plasmodium. Frontiers in cellular and infection microbiology. 2020;10:618454. doi: 10.3389/fcimb.2020.618454. PubMed PMID: 33425787; PubMed Central PMCID: PMC7793691.

2. Modrzynska K, Pfander C, Chappell L, Yu L, Suarez C, Dundas K, Gomes AR, Goulding D, Rayner JC, Choudhary J, Billker O. A Knockout Screen of ApiAP2 Genes Reveals Networks of Interacting Transcriptional Regulators Controlling the Plasmodium Life Cycle. Cell host & microbe. 2017;21(1):11–22. doi: 10.1016/j.chom.2016.12.003. PubMed PMID: 28081440; PubMed Central PMCID: PMC5241200.

3. Waldman BS, Schwarz D, Wadsworth MH, 2nd, Saeij JP, Shalek AK, Lourido S. Identification of a Master Regulator of Differentiation in Toxoplasma. Cell. 2020;180(2):359–72 e16. doi: 10.1016/j.cell.2019.12.013. PubMed PMID: 31955846; PubMed Central PMCID: PMC6978799.

4. Alvar J, Velez ID, Bern C, Herrero M, Desjeux P, Cano J, Jannin J, den Boer M, Team WHOLC. Leishmaniasis worldwide and global estimates of its incidence. PloS one. 2012;7(5):e35671. doi: 10.1371/journal.pone.0035671. PubMed PMID: 22693548; PubMed Central PMCID: PMC3365071.

5. WHO. Leishmaniasis 2023. Available from: https://www.who.int/news-room/fact-sheets/detail/leishmaniasis.

6. Maia C, Conceicao C, Pereira A, Rocha R, Ortuno M, Munoz C, Jumakanova Z, Perez-Cutillas P, Ozbel Y, Toz S, Baneth G, Monge-Maillo B, Gasimov E, Van der Stede Y, Torres G, Gossner CM, Berriatua E. The estimated distribution of autochthonous leishmaniasis by Leishmania infantum in Europe in 2005-2020. PLoS neglected tropical diseases. 2023;17(7):e0011497. doi: 10.1371/journal.pntd.0011497. PubMed PMID: 37467280; PubMed Central PMCID: PMC10389729.

7. Barbiero A, Spinicci M, Aiello A, Maruotto M, Antonello RM, Formica G, Piccica M, Isola P, Parisio EM, Nardone M, Valentini S, Mangano V, Brunelli T, Bianchi L, Bartalesi F, Costa C, Sambo M, Tumbarello M, Sani S, Fabiani S, Rossetti B, Nencioni C, Lanari A, Aquilini D, Montorzi G, Venturini E, Galli L, Rinninella G, Falcone M, Ceriegi F, Amadori F, Vincenti A, Blanc P, Vellere I, Tacconi D, Luchi S, Moneta S, Massi D, Brogi M, Voller F, Gemmi F, Rossolini GM, Cusi MG, Bruschi F, Bartoloni A, Zammarchi L. The Uprise of Human Leishmaniasis in Tuscany, Central Italy: Clinical and Epidemiological Data from a Multicenter Study. Microorganisms. 2024;12(10). doi: 10.3390/microorganisms12101963. PubMed PMID: 39458272; PubMed Central PMCID: PMC11509187.

8. Handman E. Cell biology of Leishmania. Advances in parasitology. 1999;44:1–39. doi: 10.1016/s0065-308x(08)60229-8. PubMed PMID: 10563394.

9. Clayton CE. Life without transcriptional control? From fly to man and back again. The EMBO journal. 2002;21(8):1881–8. doi: 10.1093/emboj/21.8.1881. PubMed PMID: 11953307; PubMed Central PMCID: PMC125970.

10. Liang XH, Haritan A, Uliel S, Michaeli S. trans and cis splicing in trypanosomatids: mechanism, factors, and regulation. Eukaryotic cell. 2003;2(5):830–40. doi: 10.1128/EC.2.5.830-840.2003. PubMed PMID: 14555465; PubMed Central PMCID: PMC219355.

11. McConville MJ, Ralton JE. Developmentally regulated changes in the cell surface architecture of Leishmania parasites. Behring Institute Mitteilungen. 1997(99):34–43. PubMed PMID: 9303200.

12. McConville MJ, de Souza D, Saunders E, Likic VA, Naderer T. Living in a phagolysosome; metabolism of Leishmania amastigotes. Trends in parasitology. 2007;23(8):368–75. doi: 10.1016/j.pt.2007.06.009. PubMed PMID: 17606406.

13. Rosenzweig D, Smith D, Opperdoes F, Stern S, Olafson RW, Zilberstein D. Retooling Leishmania metabolism: from sand fly gut to human macrophage. FASEB journal : official publication of the Federation of American Societies for Experimental Biology. 2008;22(2):590–602. doi: 10.1096/fj.07-9254com. PubMed PMID: 17884972.

14. McConville MJ, Naderer T. Metabolic pathways required for the intracellular survival of Leishmania. Annual review of microbiology. 2011;65:543–61. doi: 10.1146/annurev-micro-090110-102913. PubMed PMID: 21721937.

15. Sunter J, Gull K. Shape, form, function and Leishmania pathogenicity: from textbook descriptions to biological understanding. Open biology. 2017;7(9). doi: 10.1098/rsob.170165. PubMed PMID: 28903998; PubMed Central PMCID: PMC5627057.

16. Dandugudumula R, Fischer-Weinberger R, Zilberstein D. Morphogenesis Dynamics in Leishmania Differentiation. Pathogens. 2022;11(9). doi: 10.3390/pathogens11090952. PubMed PMID: 36145385; PubMed Central PMCID: PMC9505065.

17. Trenaman A, Glover L, Hutchinson S, Horn D. A post-transcriptional respiratome regulon in trypanosomes. Nucleic acids research. 2019;47(13):7063–77. doi: 10.1093/nar/gkz455. PubMed PMID: 31127277; PubMed Central PMCID: PMC6648352.

18. Boucher N, Wu Y, Dumas C, Dube M, Sereno D, Breton M, Papadopoulou B. A common mechanism of stage-regulated gene expression in Leishmania mediated by a conserved 3’-untranslated region element. The Journal of biological chemistry. 2002;277(22):19511–20. doi: 10.1074/jbc.M200500200. PubMed PMID: 11912202.

19. McNicoll F, Muller M, Cloutier S, Boilard N, Rochette A, Dube M, Papadopoulou B. Distinct 3’-untranslated region elements regulate stage-specific mRNA accumulation and translation in Leishmania. The Journal of biological chemistry. 2005;280(42):35238–46. doi: 10.1074/jbc.M507511200. PubMed PMID: 16115874.

20. Bringaud F, Muller M, Cerqueira GC, Smith M, Rochette A, El-Sayed NM, Papadopoulou B, Ghedin E. Members of a large retroposon family are determinants of post-transcriptional gene expression in Leishmania. PLoS pathogens. 2007;3(9):1291–307. doi: 10.1371/journal.ppat.0030136. PubMed PMID: 17907803; PubMed Central PMCID: PMC2323293.

21. Azizi H, Dumas C, Papadopoulou B. The Pumilio-domain protein PUF6 contributes to SIDER2 retroposon-mediated mRNA decay in Leishmania. Rna. 2017;23(12):1874–85. doi: 10.1261/rna.062950.117. PubMed PMID: 28877997; PubMed Central PMCID: PMC5689007.

22. Clayton C, Estevez A. The exosomes of trypanosomes and other protists. Advances in experimental medicine and biology. 2011;702:39–49. doi: 10.1007/978-1-4419-7841-7_4. PubMed PMID: 21713676.

23. Schwede A, Manful T, Jha BA, Helbig C, Bercovich N, Stewart M, Clayton C. The role of deadenylation in the degradation of unstable mRNAs in trypanosomes. Nucleic acids research. 2009;37(16):5511–28. doi: 10.1093/nar/gkp571. PubMed PMID: 19596809; PubMed Central PMCID: PMC2760810.

24. Kramer S. The ApaH-like phosphatase TbALPH1 is the major mRNA decapping enzyme of trypanosomes. PLoS pathogens. 2017;13(6):e1006456. doi: 10.1371/journal.ppat.1006456. PubMed PMID: 28628654; PubMed Central PMCID: PMC5491325.

25. Yoffe Y, Zuberek J, Lewdorowicz M, Zeira Z, Keasar C, Orr-Dahan I, Jankowska-Anyszka M, Stepinski J, Darzynkiewicz E, Shapira M. Cap-binding activity of an eIF4E homolog from Leishmania. Rna. 2004;10(11):1764–75. doi: 10.1261/rna.7520404. PubMed PMID: 15388875; PubMed Central PMCID: PMC1370664.

26. Yoffe Y, Zuberek J, Lerer A, Lewdorowicz M, Stepinski J, Altmann M, Darzynkiewicz E, Shapira M. Binding specificities and potential roles of isoforms of eukaryotic initiation factor 4E in Leishmania. Eukaryotic cell. 2006;5(12):1969–79. doi: 10.1128/EC.00230-06. PubMed PMID: 17041189; PubMed Central PMCID: PMC1694823.

27. David M, Gabdank I, Ben-David M, Zilka A, Orr I, Barash D, Shapira M. Preferential translation of Hsp83 in Leishmania requires a thermosensitive polypyrimidine-rich element in the 3’ UTR and involves scanning of the 5’ UTR. Rna. 2010;16(2):364–74. doi: 10.1261/rna.1874710. PubMed PMID: 20040590; PubMed Central PMCID: PMC2811665.

28. Freire ER, Sturm NR, Campbell DA, de Melo Neto OP. The Role of Cytoplasmic mRNA Cap-Binding Protein Complexes in Trypanosoma brucei and Other Trypanosomatids. Pathogens. 2017;6(4). doi: 10.3390/pathogens6040055. PubMed PMID: 29077018; PubMed Central PMCID: PMC5750579.

29. Shrivastava R, Tupperwar N, Schwartz B, Baron N, Shapira M. LeishIF4E-5 Is a Promastigote-Specific Cap-Binding Protein in Leishmania. International journal of molecular sciences. 2021;22(8). doi: 10.3390/ijms22083979. PubMed PMID: 33921489; PubMed Central PMCID: PMC8069130.

30. Assis LA, Santos Filho MVC, da Cruz Silva JR, Bezerra MJR, de Aquino I, Merlo KC, Holetz FB, Probst CM, Rezende AM, Papadopoulou B, da Costa Lima TDC, de Melo Neto OP. Identification of novel proteins and mRNAs differentially bound to the Leishmania Poly(A) Binding Proteins reveals a direct association between PABP1, the RNA-binding protein RBP23 and mRNAs encoding ribosomal proteins. PLoS neglected tropical diseases. 2021;15(10):e0009899. doi: 10.1371/journal.pntd.0009899. PubMed PMID: 34705820; PubMed Central PMCID: PMC8575317 competing interests.

31. Piel L, Rajan KS, Bussotti G, Varet H, Legendre R, Proux C, Douche T, Giai-Gianetto Q, Chaze T, Cokelaer T, Vojtkova B, Gordon-Bar N, Doniger T, Cohen-Chalamish S, Rengaraj P, Besse C, Boland A, Sadlova J, Deleuze JF, Matondo M, Unger R, Volf P, Michaeli S, Pescher P, Spath GF. Experimental evolution links post-transcriptional regulation to Leishmania fitness gain. PLoS pathogens. 2022;18(3):e1010375. doi: 10.1371/journal.ppat.1010375. PubMed PMID: 35294501; PubMed Central PMCID: PMC8959184.

32. Rodriguez-Almonacid CC, Kellogg MK, Karamyshev AL, Karamysheva ZN. Ribosome Specialization in Protozoa Parasites. International journal of molecular sciences. 2023;24(8). doi: 10.3390/ijms24087484. PubMed PMID: 37108644; PubMed Central PMCID: PMC10138883.

33. Gutierrez Guarnizo SA, Tikhonova EB, Karamyshev AL, Muskus CE, Karamysheva ZN. Translational reprogramming as a driver of antimony-drug resistance in Leishmania. Nature communications. 2023;14(1):2605. doi: 10.1038/s41467-023-38221-1. PubMed PMID: 37147291; PubMed Central PMCID: PMC10163012.

34. Rajan KS, Aryal S, Hiregange DG, Bashan A, Madmoni H, Olami M, Doniger T, Cohen-Chalamish S, Pescher P, Taoka M, Nobe Y, Fedorenko A, Bose T, Zimermann E, Prina E, Aharon-Hefetz N, Pilpel Y, Isobe T, Unger R, Spath GF, Yonath A, Michaeli S. Structural and mechanistic insights into the function of Leishmania ribosome lacking a single pseudouridine modification. Cell reports. 2024;43(5):114203. doi: 10.1016/j.celrep.2024.114203. PubMed PMID: 38722744.

35. Besteiro S, Williams RA, Morrison LS, Coombs GH, Mottram JC. Endosome sorting and autophagy are essential for differentiation and virulence of Leishmania major. The Journal of biological chemistry. 2006;281(16):11384–96. doi: 10.1074/jbc.M512307200. PubMed PMID: 16497676.

36. Williams RA, Tetley L, Mottram JC, Coombs GH. Cysteine peptidases CPA and CPB are vital for autophagy and differentiation in Leishmania mexicana. Molecular microbiology. 2006;61(3):655–74. doi: 10.1111/j.1365-2958.2006.05274.x. PubMed PMID: 16803590.

37. Besteiro S, Williams RA, Coombs GH, Mottram JC. Protein turnover and differentiation in Leishmania. International journal for parasitology. 2007;37(10):1063–75. doi: 10.1016/j.ijpara.2007.03.008. PubMed PMID: 17493624; PubMed Central PMCID: PMC2244715.

38. Williams RA, Smith TK, Cull B, Mottram JC, Coombs GH. ATG5 is essential for ATG8-dependent autophagy and mitochondrial homeostasis in Leishmania major. PLoS pathogens. 2012;8(5):e1002695. doi: 10.1371/journal.ppat.1002695. PubMed PMID: 22615560; PubMed Central PMCID: PMC3355087.

39. Williams RA, Mottram JC, Coombs GH. Distinct roles in autophagy and importance in infectivity of the two ATG4 cysteine peptidases of Leishmania major. The Journal of biological chemistry. 2013;288(5):3678–90. doi: 10.1074/jbc.M112.415372. PubMed PMID: 23166325; PubMed Central PMCID: PMC3561585.

40. Sakamoto H, Nakada-Tsukui K, Besteiro S. The Autophagy Machinery in Human-Parasitic Protists; Diverse Functions for Universally Conserved Proteins. Cells. 2021;10(5). doi: 10.3390/cells10051258. PubMed PMID: 34069694; PubMed Central PMCID: PMC8161075.

41. Casgrain PA, Martel C, McMaster WR, Mottram JC, Olivier M, Descoteaux A. Cysteine Peptidase B Regulates Leishmania mexicana Virulence through the Modulation of GP63 Expression. PLoS pathogens. 2016;12(5):e1005658. doi: 10.1371/journal.ppat.1005658. PubMed PMID: 27191844; PubMed Central PMCID: PMC4871588.

42. Pescher P, Blisnick T, Bastin P, Spath GF. Quantitative proteome profiling informs on phenotypic traits that adapt Leishmania donovani for axenic and intracellular proliferation. Cellular microbiology. 2011;13(7):978–91. doi: 10.1111/j.1462-5822.2011.01593.x. PubMed PMID: 21501362.

43. Li H, Handsaker B, Wysoker A, Fennell T, Ruan J, Homer N, Marth G, Abecasis G, Durbin R, Genome Project Data Processing S. The Sequence Alignment/Map format and SAMtools. Bioinformatics. 2009;25(16):2078–9. doi: 10.1093/bioinformatics/btp352. PubMed PMID: 19505943; PubMed Central PMCID: PMC2723002.

44. DePristo MA, Banks E, Poplin R, Garimella KV, Maguire JR, Hartl C, Philippakis AA, del Angel G, Rivas MA, Hanna M, McKenna A, Fennell TJ, Kernytsky AM, Sivachenko AY, Cibulskis K, Gabriel SB, Altshuler D, Daly MJ. A framework for variation discovery and genotyping using next-generation DNA sequencing data. Nature genetics. 2011;43(5):491–8. doi: 10.1038/ng.806. PubMed PMID: 21478889; PubMed Central PMCID: PMC3083463.

45. Quinlan AR, Hall IM. BEDTools: a flexible suite of utilities for comparing genomic features. Bioinformatics. 2010;26(6):841–2. doi: 10.1093/bioinformatics/btq033. PubMed PMID: 20110278; PubMed Central PMCID: PMC2832824.

46. Prieto Barja P, Pescher P, Bussotti G, Dumetz F, Imamura H, Kedra D, Domagalska M, Chaumeau V, Himmelbauer H, Pages M, Sterkers Y, Dujardin JC, Notredame C, Spath GF. Haplotype selection as an adaptive mechanism in the protozoan pathogen Leishmania donovani. Nature ecology & evolution. 2017;1(12):1961–9. doi: 10.1038/s41559-017-0361-x. PubMed PMID: 29109466.

47. Liao Y, Smyth GK, Shi W. featureCounts: an efficient general purpose program for assigning sequence reads to genomic features. Bioinformatics. 2014;30(7):923–30. doi: 10.1093/bioinformatics/btt656. PubMed PMID: 24227677.

48. Spath GF, Bussotti G. GIP: an open-source computational pipeline for mapping genomic instability from protists to cancer cells. Nucleic acids research. 2022;50(6):e36. doi: 10.1093/nar/gkab1237. PubMed PMID: 34928370; PubMed Central PMCID: PMC8989552.

49. McKinney W. Data Structures for Statistical Computing in Python 2010. doi: doi.org/10.25080/Majora-92bf1922-00a.

50. Reback J, jbrockmendel, McKinney W, Van Den Bossche J, Augspurger T, Cloud P, Hawkins S, gfyoung, Sinhrks, Roeschke M, Klein A, Petersen T, Tratner J, She C, Ayd W. pandas-dev/pandas: Pandas 1.3.1.

51. Hunter JD. Matplotlib: A 2D Graphics Environment. Computing in Science & Engineering. 2007;9(3):90–5. doi: 10.1109/MCSE.2007.55.

52. Waskom ML. Seaborn: statistical data visualization. . Journal of Open Source Software. 2021;6(60). doi: 10.21105/joss.03021.

53. Cokelaer T. Sequana’: a Set of Snakemake NGS pipelines. Journal of Open Source Software. 2017;2(16). doi: 10.21105/joss.00352.

54. Koster J, Rahmann S. Snakemake--a scalable bioinformatics workflow engine. Bioinformatics. 2012;28(19):2520–2. doi: 10.1093/bioinformatics/bts480. PubMed PMID: 22908215.

55. Langmead B, Salzberg SL. Fast gapped-read alignment with Bowtie 2. Nature methods. 2012;9(4):357–9. doi: 10.1038/nmeth.1923. PubMed PMID: 22388286; PubMed Central PMCID: PMC3322381.

56. Love MI, Huber W, Anders S. Moderated estimation of fold change and dispersion for RNA-seq data with DESeq2. Genome biology. 2014;15(12):550. doi: 10.1186/s13059-014-0550-8. PubMed PMID: 25516281; PubMed Central PMCID: PMC4302049.

57. Martin M. Cutadapt removes adapter sequences from high-throughput sequencing reads. EMBnetjournal. 2011;17(1). doi: 10.14806/ej.17.1.200.

58. Dobin A, Davis CA, Schlesinger F, Drenkow J, Zaleski C, Jha S, Batut P, Chaisson M, Gingeras TR. STAR: ultrafast universal RNA-seq aligner. Bioinformatics. 2013;29(1):15–21. doi: 10.1093/bioinformatics/bts635. PubMed PMID: 23104886; PubMed Central PMCID: PMC3530905.

59. Team RC. R: A Language and Environment for Statistical Computing. R Foundation for Statistical Computing 2016.

60. Benjamini YHY. Controlling the False Discovery Rate: A Practical and Powerful Approach to Multiple Testing. Journal of the Royal Statistical Society: Series B (Methodological). 1995;57(1):289–300. doi: 10.1111/j.2517-6161.1995.tb02031.x.

61. Erde J, Loo RR, Loo JA. Enhanced FASP (eFASP) to increase proteome coverage and sample recovery for quantitative proteomic experiments. Journal of proteome research. 2014;13(4):1885–95. doi: 10.1021/pr4010019. PubMed PMID: 24552128; PubMed Central PMCID: PMC3993969.

62. Matheron L, van den Toorn H, Heck AJ, Mohammed S. Characterization of biases in phosphopeptide enrichment by Ti(4+)-immobilized metal affinity chromatography and TiO2 using a massive synthetic library and human cell digests. Analytical chemistry. 2014;86(16):8312–20. doi: 10.1021/ac501803z. PubMed PMID: 25068997.

63. Creek DJ, Jankevics A, Burgess KE, Breitling R, Barrett MP. IDEOM: an Excel interface for analysis of LC-MS-based metabolomics data. Bioinformatics. 2012;28(7):1048–9. doi: 10.1093/bioinformatics/bts069. PubMed PMID: 22308147.

64. Scheltema RA, Jankevics A, Jansen RC, Swertz MA, Breitling R. PeakML/mzMatch: a file format, Java library, R library, and tool-chain for mass spectrometry data analysis. Analytical chemistry. 2011;83(7):2786–93. doi: 10.1021/ac2000994. PubMed PMID: 21401061.

65. Pettersen EF, Goddard TD, Huang CC, Meng EC, Couch GS, Croll TI, Morris JH, Ferrin TE. UCSF ChimeraX: Structure visualization for researchers, educators, and developers. Protein science : a publication of the Protein Society. 2021;30(1):70–82. doi: 10.1002/pro.3943. PubMed PMID: 32881101; PubMed Central PMCID: PMC7737788.

66. Bussotti G, Piel L, Pescher P, Domagalska MA, Rajan KS, Cohen-Chalamish S, Doniger T, Hiregange DG, Myler PJ, Unger R, Michaeli S, Spath GF. Genome instability drives epistatic adaptation in the human pathogen Leishmania. Proceedings of the National Academy of Sciences of the United States of America. 2021;118(51). doi: 10.1073/pnas.2113744118. PubMed PMID: 34903666; PubMed Central PMCID: PMC8713814.

67. Perez-Riverol Y, Csordas A, Bai J, Bernal-Llinares M, Hewapathirana S, Kundu DJ, Inuganti A, Griss J, Mayer G, Eisenacher M, Perez E, Uszkoreit J, Pfeuffer J, Sachsenberg T, Yilmaz S, Tiwary S, Cox J, Audain E, Walzer M, Jarnuczak AF, Ternent T, Brazma A, Vizcaino JA. The PRIDE database and related tools and resources in 2019: improving support for quantification data. Nucleic acids research. 2019;47(D1):D442–D50. doi: 10.1093/nar/gky1106. PubMed PMID: 30395289; PubMed Central PMCID: PMC6323896.

68. Edgar R, Domrachev M, Lash AE. Gene Expression Omnibus: NCBI gene expression and hybridization array data repository. Nucleic acids research. 2002;30(1):207–10. doi: 10.1093/nar/30.1.207. PubMed PMID: 11752295; PubMed Central PMCID: PMC99122.

69. Athar A, Fullgrabe A, George N, Iqbal H, Huerta L, Ali A, Snow C, Fonseca NA, Petryszak R, Papatheodorou I, Sarkans U, Brazma A. ArrayExpress update - from bulk to single-cell expression data. Nucleic acids research. 2019;47(D1):D711–D5. doi: 10.1093/nar/gky964. PubMed PMID: 30357387; PubMed Central PMCID: PMC6323929.

70. Bussotti G, Li B, Pescher P, Vojtkova B, Louradour I, Pruzinova K, Sadlova J, Volf P, Spath GF. Leishmania allelic selection during experimental sand fly infection correlates with mutational signatures of oxidative DNA damage. Proceedings of the National Academy of Sciences of the United States of America. 2023;120(10):e2220828120. doi: 10.1073/pnas.2220828120. PubMed PMID: 36848551; PubMed Central PMCID: PMC10013807.

71. Marques CA, Dickens NJ, Paape D, Campbell SJ, McCulloch R. Genome-wide mapping reveals single-origin chromosome replication in Leishmania, a eukaryotic microbe. Genome biology. 2015;16:230. doi: 10.1186/s13059-015-0788-9. PubMed PMID: 26481451; PubMed Central PMCID: PMC4612428.

72. Bussotti G, Gouzelou E, Cortes Boite M, Kherachi I, Harrat Z, Eddaikra N, Mottram JC, Antoniou M, Christodoulou V, Bali A, Guerfali FZ, Laouini D, Mukhtar M, Dumetz F, Dujardin JC, Smirlis D, Lechat P, Pescher P, El Hamouchi A, Lemrani M, Chicharro C, Llanes-Acevedo IP, Botana L, Cruz I, Moreno J, Jeddi F, Aoun K, Bouratbine A, Cupolillo E, Spath GF. Leishmania Genome Dynamics during Environmental Adaptation Reveal Strain-Specific Differences in Gene Copy Number Variation, Karyotype Instability, and Telomeric Amplification. mBio. 2018;9(6). doi: 10.1128/mBio.01399-18. PubMed PMID: 30401775; PubMed Central PMCID: PMC6222132.

73. Kloehn J, Saunders EC, O’Callaghan S, Dagley MJ, McConville MJ. Characterization of metabolically quiescent Leishmania parasites in murine lesions using heavy water labeling. PLoS pathogens. 2015;11(2):e1004683. doi: 10.1371/journal.ppat.1004683. PubMed PMID: 25714830; PubMed Central PMCID: PMC4340956.

74. Bussotti G, Benkahla A, Jeddi F, Souiai O, Aoun K, Spath GF, Bouratbine A. Nuclear and mitochondrial genome sequencing of North-African Leishmania infantum isolates from cured and relapsed visceral leishmaniasis patients reveals variations correlating with geography and phenotype. Microbial genomics. 2020;6(10). doi: 10.1099/mgen.0.000444. PubMed PMID: 32975503; PubMed Central PMCID: PMC7660250.

75. Cuypers B, Meysman P, Erb I, Bittremieux W, Valkenborg D, Baggerman G, Mertens I, Sundar S, Khanal B, Notredame C, Dujardin JC, Domagalska MA, Laukens K. Four layer multi-omics reveals molecular responses to aneuploidy in Leishmania. PLoS pathogens. 2022;18(9):e1010848. doi: 10.1371/journal.ppat.1010848. PubMed PMID: 36149920; PubMed Central PMCID: PMC9534393.

76. Dumetz F, Imamura H, Sanders M, Seblova V, Myskova J, Pescher P, Vanaerschot M, Meehan CJ, Cuypers B, De Muylder G, Spath GF, Bussotti G, Vermeesch JR, Berriman M, Cotton JA, Volf P, Dujardin JC, Domagalska MA. Modulation of Aneuploidy in Leishmania donovani during Adaptation to Different In Vitro and In Vivo Environments and Its Impact on Gene Expression. mBio. 2017;8(3). doi: 10.1128/mBio.00599-17. PubMed PMID: 28536289; PubMed Central PMCID: PMC5442457.

77. Fiebig M, Kelly S, Gluenz E. Comparative Life Cycle Transcriptomics Revises Leishmania mexicana Genome Annotation and Links a Chromosome Duplication with Parasitism of Vertebrates. PLoS pathogens. 2015;11(10):e1005186. doi: 10.1371/journal.ppat.1005186. PubMed PMID: 26452044; PubMed Central PMCID: PMC4599935.

78. Wu Y, El Fakhry Y, Sereno D, Tamar S, Papadopoulou B. A new developmentally regulated gene family in Leishmania amastigotes encoding a homolog of amastin surface proteins. Molecular and biochemical parasitology. 2000;110(2):345–57. doi: 10.1016/s0166-6851(00)00290-5. PubMed PMID: 11071288.

79. Haile S, Dupe A, Papadopoulou B. Deadenylation-independent stage-specific mRNA degradation in Leishmania. Nucleic acids research. 2008;36(5):1634–44. doi: 10.1093/nar/gkn019. PubMed PMID: 18250085; PubMed Central PMCID: PMC2275140.

80. Saunders EC, Sernee MF, Ralton JE, McConville MJ. Metabolic stringent response in intracellular stages of Leishmania. Current opinion in microbiology. 2021;63:126–32. doi: 10.1016/j.mib.2021.07.007. PubMed PMID: 34340099.

81. Sernee MF, Ralton JE, Nero TL, Sobala LF, Kloehn J, Vieira-Lara MA, Cobbold SA, Stanton L, Pires DEV, Hanssen E, Males A, Ward T, Bastidas LM, van der Peet PL, Parker MW, Ascher DB, Williams SJ, Davies GJ, McConville MJ. A Family of Dual-Activity Glycosyltransferase-Phosphorylases Mediates Mannogen Turnover and Virulence in Leishmania Parasites. Cell host & microbe. 2019;26(3):385–99 e9. doi: 10.1016/j.chom.2019.08.009. PubMed PMID: 31513773.

82. Zhang H, Lin G. Microbial proteasomes as drug targets. PLoS pathogens. 2021;17(12):e1010058. doi: 10.1371/journal.ppat.1010058. PubMed PMID: 34882737; PubMed Central PMCID: PMC8659679.

83. Omura S, Crump A. Lactacystin: first-in-class proteasome inhibitor still excelling and an exemplar for future antibiotic research. The Journal of antibiotics. 2019;72(4):189–201. doi: 10.1038/s41429-019-0141-8. PubMed PMID: 30755736; PubMed Central PMCID: PMC6760633.

84. Nagle A, Biggart A, Be C, Srinivas H, Hein A, Caridha D, Sciotti RJ, Pybus B, Kreishman-Deitrick M, Bursulaya B, Lai YH, Gao MY, Liang F, Mathison CJN, Liu X, Yeh V, Smith J, Lerario I, Xie Y, Chianelli D, Gibney M, Berman A, Chen YL, Jiricek J, Davis LC, Liu X, Ballard J, Khare S, Eggimann FK, Luneau A, Groessl T, Shapiro M, Richmond W, Johnson K, Rudewicz PJ, Rao SPS, Thompson C, Tuntland T, Spraggon G, Glynne RJ, Supek F, Wiesmann C, Molteni V. Discovery and Characterization of Clinical Candidate LXE408 as a Kinetoplastid-Selective Proteasome Inhibitor for the Treatment of Leishmaniases. Journal of medicinal chemistry. 2020;63(19):10773–81. doi: 10.1021/acs.jmedchem.0c00499. PubMed PMID: 32667203; PubMed Central PMCID: PMC7549094 authors are employees or former employees of GNF or NITD or NIBR: A.S.N., A.B., C.B., S.H., A.H., B.B., Y.H.L., M.-Y.G., F.L., C.J.N.M., X.L., V.Y., J.S., I.L., Y.X., D.C., M.G., A.B., Y.-L.C., J.J., L.C.D., X.L., J.B., S.K., F.K.E., A.L., T.G., M.S., W.R., K.J., P.J.R., S.P.S.R., T.T., G.S., R.J.G., F.S., C.W., and V.M.

85. Silva-Jardim I, Horta MF, Ramalho-Pinto FJ. The Leishmania chagasi proteasome: role in promastigotes growth and amastigotes survival within murine macrophages. Acta tropica. 2004;91(2):121–30. doi: 10.1016/j.actatropica.2004.03.007. PubMed PMID: 15234661.

86. Kumar P, Sundar S, Singh N. Degradation of pteridine reductase 1 (PTR1) enzyme during growth phase in the protozoan parasite Leishmania donovani. Experimental parasitology. 2007;116(2):182–9. doi: 10.1016/j.exppara.2006.12.008. PubMed PMID: 17275814.

87. Perez-Pertejo Y, Alvarez-Velilla R, Estrada CG, Balana-Fouce R, Reguera RM. Leishmania donovani: proteasome-mediated down-regulation of methionine adenosyltransferase. Parasitology. 2011;138(9):1082–92. doi: 10.1017/S0031182011000862. PubMed PMID: 21813028.

88. Vince JE, Tull D, Landfear S, McConville MJ. Lysosomal degradation of Leishmania hexose and inositol transporters is regulated in a stage-, nutrient- and ubiquitin-dependent manner. International journal for parasitology. 2011;41(7):791–800. doi: 10.1016/j.ijpara.2011.02.003. PubMed PMID: 21447343; PubMed Central PMCID: PMC3592973.

89. Khare S, Nagle AS, Biggart A, Lai YH, Liang F, Davis LC, Barnes SW, Mathison CJ, Myburgh E, Gao MY, Gillespie JR, Liu X, Tan JL, Stinson M, Rivera IC, Ballard J, Yeh V, Groessl T, Federe G, Koh HX, Venable JD, Bursulaya B, Shapiro M, Mishra PK, Spraggon G, Brock A, Mottram JC, Buckner FS, Rao SP, Wen BG, Walker JR, Tuntland T, Molteni V, Glynne RJ, Supek F. Proteasome inhibition for treatment of leishmaniasis, Chagas disease and sleeping sickness. Nature. 2016;537(7619):229–33. doi: 10.1038/nature19339. PubMed PMID: 27501246; PubMed Central PMCID: PMC5161665.

90. Wyllie S, Brand S, Thomas M, De Rycker M, Chung CW, Pena I, Bingham RP, Bueren-Calabuig JA, Cantizani J, Cebrian D, Craggs PD, Ferguson L, Goswami P, Hobrath J, Howe J, Jeacock L, Ko EJ, Korczynska J, MacLean L, Manthri S, Martinez MS, Mata-Cantero L, Moniz S, Nuhs A, Osuna-Cabello M, Pinto E, Riley J, Robinson S, Rowland P, Simeons FRC, Shishikura Y, Spinks D, Stojanovski L, Thomas J, Thompson S, Viayna Gaza E, Wall RJ, Zuccotto F, Horn D, Ferguson MAJ, Fairlamb AH, Fiandor JM, Martin J, Gray DW, Miles TJ, Gilbert IH, Read KD, Marco M, Wyatt PG. Preclinical candidate for the treatment of visceral leishmaniasis that acts through proteasome inhibition. Proceedings of the National Academy of Sciences of the United States of America. 2019;116(19):9318–23. doi: 10.1073/pnas.1820175116. PubMed PMID: 30962368; PubMed Central PMCID: PMC6511062.

91. Burge RJ, Damianou A, Wilkinson AJ, Rodenko B, Mottram JC. Leishmania differentiation requires ubiquitin conjugation mediated by a UBC2-UEV1 E2 complex. PLoS pathogens. 2020;16(10):e1008784. doi: 10.1371/journal.ppat.1008784. PubMed PMID: 33108402; PubMed Central PMCID: PMC7647121.

92. Damianou A, Burge RJ, Catta-Preta CMC, Geoghegan V, Nievas YR, Newling K, Brown E, Burchmore R, Rodenko B, Mottram JC. Essential roles for deubiquitination in Leishmania life cycle progression. PLoS pathogens. 2020;16(6):e1008455. doi: 10.1371/journal.ppat.1008455. PubMed PMID: 32544189; PubMed Central PMCID: PMC7319358.

93. Parsons M, Worthey EA, Ward PN, Mottram JC. Comparative analysis of the kinomes of three pathogenic trypanosomatids: Leishmania major, Trypanosoma brucei and Trypanosoma cruzi. BMC genomics. 2005;6:127. doi: 10.1186/1471-2164-6-127. PubMed PMID: 16164760; PubMed Central PMCID: PMC1266030.

94. Morales MA, Renaud O, Faigle W, Shorte SL, Spath GF. Over-expression of Leishmania major MAP kinases reveals stage-specific induction of phosphotransferase activity. International journal for parasitology. 2007;37(11):1187–99. doi: 10.1016/j.ijpara.2007.03.006. PubMed PMID: 17481635.

95. Rotureau B, Morales MA, Bastin P, Spath GF. The flagellum-mitogen-activated protein kinase connection in Trypanosomatids: a key sensory role in parasite signalling and development? Cellular microbiology. 2009;11(5):710–8. doi: 10.1111/j.1462-5822.2009.01295.x. PubMed PMID: 19207727.

96. Morales MA, Pescher P, Spath GF. Leishmania major MPK7 protein kinase activity inhibits intracellular growth of the pathogenic amastigote stage. Eukaryotic cell. 2010;9(1):22–30. doi: 10.1128/EC.00196-09. PubMed PMID: 19801421; PubMed Central PMCID: PMC2805286.

97. Potenza M, Schenkman S, Laverriere M, Tellez-Inon MT. Functional characterization of TcCYC2 cyclin from Trypanosoma cruzi. Experimental parasitology. 2012;132(4):537–45. doi: 10.1016/j.exppara.2012.09.002. PubMed PMID: 22982808.

98. Dacher M, Morales MA, Pescher P, Leclercq O, Rachidi N, Prina E, Cayla M, Descoteaux A, Spath GF. Probing druggability and biological function of essential proteins in Leishmania combining facilitated null mutant and plasmid shuffle analyses. Molecular microbiology. 2014;93(1):146–66. doi: 10.1111/mmi.12648. PubMed PMID: 24823804.

99. Cayla M, Rachidi N, Leclercq O, Schmidt-Arras D, Rosenqvist H, Wiese M, Spath GF. Transgenic analysis of the Leishmania MAP kinase MPK10 reveals an auto-inhibitory mechanism crucial for stage-regulated activity and parasite viability. PLoS pathogens. 2014;10(9):e1004347. doi: 10.1371/journal.ppat.1004347. PubMed PMID: 25232945; PubMed Central PMCID: PMC4169501.

100. Jones NG, Thomas EB, Brown E, Dickens NJ, Hammarton TC, Mottram JC. Regulators of Trypanosoma brucei cell cycle progression and differentiation identified using a kinome-wide RNAi screen. PLoS pathogens. 2014;10(1):e1003886. doi: 10.1371/journal.ppat.1003886. PubMed PMID: 24453978; PubMed Central PMCID: PMC3894213.

101. Rachidi N, Taly JF, Durieu E, Leclercq O, Aulner N, Prina E, Pescher P, Notredame C, Meijer L, Spath GF. Pharmacological assessment defines Leishmania donovani casein kinase 1 as a drug target and reveals important functions in parasite viability and intracellular infection. Antimicrobial agents and chemotherapy. 2014;58(3):1501–15. doi: 10.1128/AAC.02022-13. PubMed PMID: 24366737; PubMed Central PMCID: PMC3957854.

102. Baker N, Catta-Preta CMC, Neish R, Sadlova J, Powell B, Alves-Ferreira EVC, Geoghegan V, Carnielli JBT, Newling K, Hughes C, Vojtkova B, Anand J, Mihut A, Walrad PB, Wilson LG, Pitchford JW, Volf P, Mottram JC. Systematic functional analysis of Leishmania protein kinases identifies regulators of differentiation or survival. Nature communications. 2021;12(1):1244. doi: 10.1038/s41467-021-21360-8. PubMed PMID: 33623024; PubMed Central PMCID: PMC7902614.

103. Cayla M, Nievas YR, Matthews KR, Mottram JC. Distinguishing functions of trypanosomatid protein kinases. Trends in parasitology. 2022;38(11):950–61. doi: 10.1016/j.pt.2022.08.009. PubMed PMID: 36075845.

104. Grunebast J, Lorenzen S, Clos J. Genome-wide quantification of polycistronic transcription in Leishmania major. mBio. 2025;16(1):e0224124. doi: 10.1128/mbio.02241-24. PubMed PMID: 39584812; PubMed Central PMCID: PMC11708010.

105. Muller-McNicoll M, Rossbach O, Hui J, Medenbach J. Auto-regulatory feedback by RNA-binding proteins. Journal of molecular cell biology. 2019;11(10):930–9. doi: 10.1093/jmcb/mjz043. PubMed PMID: 31152582; PubMed Central PMCID: PMC6884704.

106. Perrimon N, Pitsouli C, Shilo BZ. Signaling mechanisms controlling cell fate and embryonic patterning. Cold Spring Harbor perspectives in biology. 2012;4(8):a005975. doi: 10.1101/cshperspect.a005975. PubMed PMID: 22855721; PubMed Central PMCID: PMC3405863.

107. Erben E, Leiss K, Liu B, Gil DI, Helbig C, Clayton C. Insights into the functions and RNA binding of Trypanosoma brucei ZC3H22, RBP9 and DRBD7. Parasitology. 2021;148(10):1186–95. doi: 10.1017/S0031182021000123. PubMed PMID: 33536101; PubMed Central PMCID: PMC8312216.

108. Walrad PB, Capewell P, Fenn K, Matthews KR. The post-transcriptional trans-acting regulator, TbZFP3, co-ordinates transmission-stage enriched mRNAs in Trypanosoma brucei. Nucleic acids research. 2012;40(7):2869–83. doi: 10.1093/nar/gkr1106. PubMed PMID: 22140102; PubMed Central PMCID: PMC3326296.

109. Mugo E, Clayton C. Expression of the RNA-binding protein RBP10 promotes the bloodstream-form differentiation state in Trypanosoma brucei. PLoS pathogens. 2017;13(8):e1006560. doi: 10.1371/journal.ppat.1006560. PubMed PMID: 28800584; PubMed Central PMCID: PMC5568443.

110. Avila CC, Mule SN, Rosa-Fernandes L, Viner R, Barison MJ, Costa-Martins AG, Oliveira GS, Teixeira MMG, Marinho CRF, Silber AM, Palmisano G. Proteome-Wide Analysis of Trypanosoma cruzi Exponential and Stationary Growth Phases Reveals a Subcellular Compartment-Specific Regulation. Genes. 2018;9(8). doi: 10.3390/genes9080413. PubMed PMID: 30111733; PubMed Central PMCID: PMC6115888.

111. Fritz M, Vanselow J, Sauer N, Lamer S, Goos C, Siegel TN, Subota I, Schlosser A, Carrington M, Kramer S. Novel insights into RNP granules by employing the trypanosome’s microtubule skeleton as a molecular sieve. Nucleic acids research. 2015;43(16):8013–32. doi: 10.1093/nar/gkv731. PubMed PMID: 26187993; PubMed Central PMCID: PMC4652759.

112. Cargnello M, Roux PP. Activation and function of the MAPKs and their substrates, the MAPK-activated protein kinases. Microbiology and molecular biology reviews : MMBR. 2011;75(1):50–83. doi: 10.1128/MMBR.00031-10. PubMed PMID: 21372320; PubMed Central PMCID: PMC3063353.

113. Erdmann M, Scholz A, Melzer IM, Schmetz C, Wiese M. Interacting protein kinases involved in the regulation of flagellar length. Molecular biology of the cell. 2006;17(4):2035–45. doi: 10.1091/mbc.e05-10-0976. PubMed PMID: 16467378; PubMed Central PMCID: PMC1415332.

114. Johnson JL, Yaron TM, Huntsman EM, Kerelsky A, Song J, Regev A, Lin TY, Liberatore K, Cizin DM, Cohen BM, Vasan N, Ma Y, Krismer K, Robles JT, van de Kooij B, van Vlimmeren AE, Andree-Busch N, Kaufer NF, Dorovkov MV, Ryazanov AG, Takagi Y, Kastenhuber ER, Goncalves MD, Hopkins BD, Elemento O, Taatjes DJ, Maucuer A, Yamashita A, Degterev A, Uduman M, Lu J, Landry SD, Zhang B, Cossentino I, Linding R, Blenis J, Hornbeck PV, Turk BE, Yaffe MB, Cantley LC. An atlas of substrate specificities for the human serine/threonine kinome. Nature. 2023;613(7945):759–66. doi: 10.1038/s41586-022-05575-3. PubMed PMID: 36631611; PubMed Central PMCID: PMC9876800 and is a founder and receives research support from Petra Pharmaceuticals; is listed as an inventor on a patent (WO2019232403A1, Weill Cornell Medicine) for combination therapy for PI3K-associated disease or disorder, and the identification of therapeutic interventions to improve response to PI3K inhibitors for cancer treatment; is a co-founder and shareholder in Faeth Therapeutics; has equity in and consults for Cell Signaling Technologies, Volastra, Larkspur and 1 Base Pharmaceuticals; and consults for Loxo-Lilly. M.B.Y receives research support from Cardiff Oncology. T.M.Y. is a co-founder and stockholder and is on the board of directors of DESTROKE, an early-stage start-up developing mobile technology for automated clinical stroke detection. J.L.J has received consulting fees from Scorpion Therapeutics and Volastra Therapeutics. O.E. is a founder and equity holder of Volastra Therapeutics and OneThree Biotech; is a member of the scientific advisory board of Owkin, Freenome, Genetic Intelligence, Acuamark and Champions Oncology; and receives research support from Eli Lilly, Janssen and Sanofi. D.J.T. is a member of the scientific advisory board at Dewpoint Therapeutics. A.D. is an equity holder of Denali Therapeutics; and receives research support from Interline Therapeutics. N.V. reports consulting activities for Novartis and is on the scientific advisory board of Heligenics. M.D.G. is a co-founder and shareholder of Faeth Therapeutics, which is developing dietary and pharmacological therapies for cancer; and has received speaking and/or consulting fees from Pfizer, Novartis, Scorpion Therapeutics and Faeth Therapeutics.

115. Ekerot M, Stavridis MP, Delavaine L, Mitchell MP, Staples C, Owens DM, Keenan ID, Dickinson RJ, Storey KG, Keyse SM. Negative-feedback regulation of FGF signalling by DUSP6/MKP-3 is driven by ERK1/2 and mediated by Ets factor binding to a conserved site within the DUSP6/MKP-3 gene promoter. The Biochemical journal. 2008;412(2):287–98. doi: 10.1042/BJ20071512. PubMed PMID: 18321244; PubMed Central PMCID: PMC2474557.

116. Hoffmann I, Clarke PR, Marcote MJ, Karsenti E, Draetta G. Phosphorylation and activation of human cdc25-C by cdc2--cyclin B and its involvement in the self-amplification of MPF at mitosis. The EMBO journal. 1993;12(1):53–63. doi: 10.1002/j.1460-2075.1993.tb05631.x. PubMed PMID: 8428594; PubMed Central PMCID: PMC413175.

117. Zinoviev A, Leger M, Wagner G, Shapira M. A novel 4E-interacting protein in Leishmania is involved in stage-specific translation pathways. Nucleic acids research. 2011;39(19):8404–15. doi: 10.1093/nar/gkr555. PubMed PMID: 21764780; PubMed Central PMCID: PMC3201875.

118. Shrivastava R, Tupperwar N, Drory-Retwitzer M, Shapira M. Deletion of a Single LeishIF4E-3 Allele by the CRISPR-Cas9 System Alters Cell Morphology and Infectivity of Leishmania. mSphere. 2019;4(5). doi: 10.1128/mSphere.00450-19. PubMed PMID: 31484740; PubMed Central PMCID: PMC6731530.

119. Tupperwar N, Shrivastava R, Shapira M. LeishIF4E1 Deletion Affects the Promastigote Proteome, Morphology, and Infectivity. mSphere. 2019;4(6). doi: 10.1128/mSphere.00625-19. PubMed PMID: 31722993; PubMed Central PMCID: PMC6854042.

120. Tupperwar N, Shrivastava R, Baron N, Korchev O, Dahan I, Shapira M. Characterization of an Atypical eIF4E Ortholog in Leishmania, LeishIF4E-6. International journal of molecular sciences. 2021;22(23). doi: 10.3390/ijms222312720. PubMed PMID: 34884522; PubMed Central PMCID: PMC8657474.

121. Baron N, Purushotham R, Pullaiahgari D, Bose P, Zarivach R, Shapira M. LeishIF4E2 is a cap-binding protein that plays a role in Leishmania cell cycle progression. FASEB journal: official publication of the Federation of American Societies for Experimental Biology. 2024;38(1):e23367. doi: 10.1096/fj.202301665R. PubMed PMID: 38095329.

122. Baird TD, Wek RC. Eukaryotic initiation factor 2 phosphorylation and translational control in metabolism. Advances in nutrition. 2012;3(3):307–21. doi: 10.3945/an.112.002113. PubMed PMID: 22585904; PubMed Central PMCID: PMC3649462.

123. Sloan KE, Warda AS, Sharma S, Entian KD, Lafontaine DLJ, Bohnsack MT. Tuning the ribosome: The influence of rRNA modification on eukaryotic ribosome biogenesis and function. RNA biology. 2017;14(9):1138–52. doi: 10.1080/15476286.2016.1259781. PubMed PMID: 27911188; PubMed Central PMCID: PMC5699541.

124. Imami K, Milek M, Bogdanow B, Yasuda T, Kastelic N, Zauber H, Ishihama Y, Landthaler M, Selbach M. Phosphorylation of the Ribosomal Protein RPL12/uL11 Affects Translation during Mitosis. Molecular cell. 2018;72(1):84–98 e9. doi: 10.1016/j.molcel.2018.08.019. PubMed PMID: 30220558.

125. Merrick WC, Pavitt GD. Protein Synthesis Initiation in Eukaryotic Cells. Cold Spring Harbor perspectives in biology. 2018;10(12). doi: 10.1101/cshperspect.a033092. PubMed PMID: 29735639; PubMed Central PMCID: PMC6280705.

126. Piazzi M, Bavelloni A, Gallo A, Faenza I, Blalock WL. Signal Transduction in Ribosome Biogenesis: A Recipe to Avoid Disaster. International journal of molecular sciences. 2019;20(11). doi: 10.3390/ijms20112718. PubMed PMID: 31163577; PubMed Central PMCID: PMC6600399.

127. Jungers CF, Elliff JM, Masson-Meyers DS, Phiel CJ, Origanti S. Regulation of eukaryotic translation initiation factor 6 dynamics through multisite phosphorylation by GSK3. The Journal of biological chemistry. 2020;295(36):12796–813. doi: 10.1074/jbc.RA120.013324. PubMed PMID: 32703900; PubMed Central PMCID: PMC7476728.

128. Janin M, Coll-SanMartin L, Esteller M. Disruption of the RNA modifications that target the ribosome translation machinery in human cancer. Molecular cancer. 2020;19(1):70. doi: 10.1186/s12943-020-01192-8. PubMed PMID: 32241281; PubMed Central PMCID: PMC7114786.

129. Bohlen J, Roiuk M, Teleman AA. Phosphorylation of ribosomal protein S6 differentially affects mRNA translation based on ORF length. Nucleic acids research. 2021;49(22):13062–74. doi: 10.1093/nar/gkab1157. PubMed PMID: 34871442; PubMed Central PMCID: PMC8682771.

130. Khoshnevis S, Dreggors-Walker RE, Marchand V, Motorin Y, Ghalei H. Ribosomal RNA 2’-O-methylations regulate translation by impacting ribosome dynamics. Proceedings of the National Academy of Sciences of the United States of America. 2022;119(12):e2117334119. doi: 10.1073/pnas.2117334119. PubMed PMID: 35294285; PubMed Central PMCID: PMC8944910.

131. Das T, Shin SC, Song EJ, Kim EE. Regulation of Deubiquitinating Enzymes by Post-Translational Modifications. International journal of molecular sciences. 2020;21(11). doi: 10.3390/ijms21114028. PubMed PMID: 32512887; PubMed Central PMCID: PMC7312083.

132. Wang F, Ning S, Yu B, Wang Y. USP14: Structure, Function, and Target Inhibition. Frontiers in pharmacology. 2021;12:801328. doi: 10.3389/fphar.2021.801328. PubMed PMID: 35069211; PubMed Central PMCID: PMC8766727.

133. Gonzalez J, Ramalho-Pinto FJ, Frevert U, Ghiso J, Tomlinson S, Scharfstein J, Corey EJ, Nussenzweig V. Proteasome activity is required for the stage-specific transformation of a protozoan parasite. The Journal of experimental medicine. 1996;184(5):1909–18. doi: 10.1084/jem.184.5.1909. PubMed PMID: 8920878; PubMed Central PMCID: PMC2192890.

134. Cardoso J, Soares MJ, Menna-Barreto RF, Le Bloas R, Sotomaior V, Goldenberg S, Krieger MA. Inhibition of proteasome activity blocks Trypanosoma cruzi growth and metacyclogenesis. Parasitology research. 2008;103(4):941–51. doi: 10.1007/s00436-008-1081-6. PubMed PMID: 18581141.

135. Cardoso J, Lima Cde P, Leal T, Gradia DF, Fragoso SP, Goldenberg S, De Sa RG, Krieger MA. Analysis of proteasomal proteolysis during the in vitro metacyclogenesis of Trypanosoma cruzi. PloS one. 2011;6(6):e21027. doi: 10.1371/journal.pone.0021027. PubMed PMID: 21698116; PubMed Central PMCID: PMC3117861.

136. Munoz C, San Francisco J, Gutierrez B, Gonzalez J. Role of the Ubiquitin-Proteasome Systems in the Biology and Virulence of Protozoan Parasites. BioMed research international. 2015;2015:141526. doi: 10.1155/2015/141526. PubMed PMID: 26090380; PubMed Central PMCID: PMC4452248.

137. Bijlmakers MJ. Ubiquitination and the Proteasome as Drug Targets in Trypanosomatid Diseases. Frontiers in chemistry. 2020;8:630888. doi: 10.3389/fchem.2020.630888. PubMed PMID: 33732684; PubMed Central PMCID: PMC7958763.

138. Wiese M. Leishmania MAP kinases--familiar proteins in an unusual context. International journal for parasitology. 2007;37(10):1053–62. doi: 10.1016/j.ijpara.2007.04.008. PubMed PMID: 17548090.

139. Hunter T. The age of crosstalk: phosphorylation, ubiquitination, and beyond. Molecular cell. 2007;28(5):730–8. doi: 10.1016/j.molcel.2007.11.019. PubMed PMID: 18082598.

140. Nguyen LK, Kolch W, Kholodenko BN. When ubiquitination meets phosphorylation: a systems biology perspective of EGFR/MAPK signalling. Cell communication and signaling: CCS. 2013;11:52. doi: 10.1186/1478-811X-11-52. PubMed PMID: 23902637; PubMed Central PMCID: PMC3734146.

141. Barbour H, Nkwe NS, Estavoyer B, Messmer C, Gushul-Leclaire M, Villot R, Uriarte M, Boulay K, Hlayhel S, Farhat B, Milot E, Mallette FA, Daou S, Affar EB. An inventory of crosstalk between ubiquitination and other post-translational modifications in orchestrating cellular processes. iScience. 2023;26(5):106276. doi: 10.1016/j.isci.2023.106276. PubMed PMID: 37168555; PubMed Central PMCID: PMC10165199.

142. Waddington CH. Canalization of development and genetic assimilation of acquired characters. Nature. 1959;183(4676):1654–5. doi: 10.1038/1831654a0. PubMed PMID: 13666847.

143. Cox J, Mann M. MaxQuant enables high peptide identification rates, individualized p.p.b.-range mass accuracies and proteome-wide protein quantification. Nature biotechnology. 2008;26(12):1367–72. doi: 10.1038/nbt.1511. PubMed PMID: 19029910.

144. Cox J, Neuhauser N, Michalski A, Scheltema RA, Olsen JV, Mann M. Andromeda: a peptide search engine integrated into the MaxQuant environment. Journal of proteome research. 2011;10(4):1794–805. doi: 10.1021/pr101065j. PubMed PMID: 21254760.

145. Cox J, Hein MY, Luber CA, Paron I, Nagaraj N, Mann M. Accurate proteome-wide label-free quantification by delayed normalization and maximal peptide ratio extraction, termed MaxLFQ. Molecular & cellular proteomics : MCP. 2014;13(9):2513–26. doi: 10.1074/mcp.M113.031591. PubMed PMID: 24942700; PubMed Central PMCID: PMC4159666.

146. Garcia-Garcia T, Douche T, Giai Gianetto Q, Poncet S, El Omrani N, Smits WK, Cuenot E, Matondo M, Martin-Verstraete I. In-Depth Characterization of the Clostridioides difficile Phosphoproteome to Identify Ser/Thr Kinase Substrates. Molecular & cellular proteomics : MCP. 2022;21(11):100428. doi: 10.1016/j.mcpro.2022.100428. PubMed PMID: 36252736; PubMed Central PMCID: PMC9674922.

147. Wieczorek S, Combes F, Lazar C, Giai Gianetto Q, Gatto L, Dorffer A, Hesse AM, Coute Y, Ferro M, Bruley C, Burger T. DAPAR & ProStaR: software to perform statistical analyses in quantitative discovery proteomics. Bioinformatics. 2017;33(1):135–6. doi: 10.1093/bioinformatics/btw580. PubMed PMID: 27605098; PubMed Central PMCID: PMC5408771.

148. Ritchie ME, Phipson B, Wu D, Hu Y, Law CW, Shi W, Smyth GK. limma powers differential expression analyses for RNA-sequencing and microarray studies. Nucleic acids research. 2015;43(7):e47. doi: 10.1093/nar/gkv007. PubMed PMID: 25605792; PubMed Central PMCID: PMC4402510.

149. Smyth GK. Limma: linear models for microarray data. Gentleman RC, Carey VJ, Dudoit S, Irizarry RA, Huber W, editors. New York: Springer; 2005. 397–420 p.

150. Giai Gianetto Q, Combes F, Ramus C, Bruley C, Coute Y, Burger T. Calibration plot for proteomics: A graphical tool to visually check the assumptions underlying FDR control in quantitative experiments. Proteomics. 2016;16(1):29–32. doi: 10.1002/pmic.201500189. PubMed PMID: 26572953.

151. Pounds S, Cheng C. Robust estimation of the false discovery rate. Bioinformatics. 2006;22(16):1979–87. doi: 10.1093/bioinformatics/btl328. PubMed PMID: 16777905.

152. Schwanhausser B, Busse D, Li N, Dittmar G, Schuchhardt J, Wolf J, Chen W, Selbach M. Global quantification of mammalian gene expression control. Nature. 2011;473(7347):337–42. doi: 10.1038/nature10098. PubMed PMID: 21593866.

153. Giai Gianetto Q. Statistical Analysis of Post-Translational Modifications Quantified by Label-Free Proteomics Across Multiple Biological Conditions with R: Illustration from SARS-CoV-2 Infected Cells. Methods in molecular biology. 2023;2426:267–302. doi: 10.1007/978-1-0716-1967-4_12. PubMed PMID: 36308693.

154. Gianetto QG, Wieczorek S, Couté Y, Burger T. A peptide-level multiple imputation strategy accounting for the different natures of missing values in proteomics data. bioRxiv. 2020:2020.05.29.122770. doi: 10.1101/2020.05.29.122770.

