## Supplementary data for "Five-layer systems analysis of *Leishmania* stage differentiation reveals an essential role for protein degradation in parasite development"

**Supplementary data: Materiel and method**

**Label-free, quantitative total proteome analyses.** Untreated amastigotes (ama and ama-18h) and promastigotes (pro), and lactacystin-treated parasites (ama lacta and pro lacta) were recovered and washed three times with cold PBS at 2,000 x *g* or 1,600 x *g* for 10 min at 4°C. Parasite lysates were prepared in eFASP lysis buffer (4% SDS / 0,2 % DCA / 50mM TCEP / 50 mM ammonium bicarbonate buffer pH 8) (1). Filter units and collection tubes were incubated overnight in 5% (v/v) TWEEN−20®. All buffer exchanges were carried out by centrifugation at 14 000 x *g* for 10 min. For each sample, 50 µg of proteins were transferred into 30,000 Dalton MWCO centrifugal units (Amicon® Centrifugal Filters, Merck) adjusted to 250 µl with exchange buffer (8 M urea, 0.2% DCA, 100 mM ammonium bicarbonate pH 8) and washed three times with 200 µl exchange buffer, after which samples were incubated in alkylation buffer (50 mM chloroacetamide, Urea 8M, 100 mM ammonium bicarbonate pH 8) in the dark for 1h. The alkylating agent was then replaced with 200 µl exchange buffer, followed by three washes in 200 µl digestion buffer (0,2 % DCA / 50 mM ammonium bicarbonate buffer pH 8). Digestion was performed at 37°C overnight in 100 µl with Sequencing Grade Modified Trypsin (Promega - V5111) at a Protein:Trypsin ratio 50:1. Two rounds of 50 µl of 50 mM ammonium bicarbonate pH 8 were used to recover the peptide-rich solution by centrifugation. DCA was removed by acidification and phase transfer. Finally, peptides were speed-vac dried and resuspended in 2% acetonitrile (ACN), 0.1% formic acid (FA) prior to LC-MS/MS analysis.

Tryptic peptides from eFASP digestion were analyzed on a Q Exactive Plus instrument (Thermo Fisher Scientific) coupled with an EASY nLC 1200 chromatography system (Thermo Fisher Scientific). Samples were loaded into a home-made 44 cm C18 column (1.9 µm particles, 100 Å pore size, ReproSil-Pur Basic C18, Dr. Maisch GmbH, Ammerbuch-Entringen, Germany). Column equilibration and peptide loading were performed at 900 bars in buffer A (0.1% FA). Peptides were eluted at a flow rate of 250 nl/min over 182min with a multi-step gradient from 2 to 7 % buffer B (80 % ACN, 0.1 % FA) during 5 min, 7 to 23 % buffer B for 130 min, 23 to 45 % buffer B for 20 min, 45 to 95 % buffer B for 5 min. Column temperature was set to 60°C. MS data were acquired using Xcalibur software using a data-dependent method. MS scans were acquired at a resolution of 70,000 and MS/MS scans (fixed first mass 100 m/z) at a resolution of 17,500. The AGC target and maximum injection time for the survey scans and the MS/MS scans were set to 3 x $10^6$ for 20ms and $10^6$ for 60ms, respectively. An automatic selection of the 10 most intense precursor ions was activated (Top 10) with a 40 s dynamic exclusion. The isolation window was set to 1.6 m/z and normalized collision energy fixed to 28 for HCD fragmentation. We used a minimum AGC target of $10^4$ corresponding to an intensity threshold of 1.7 x $10^5$. Unassigned precursor ion charge states as well as 1, 7, 8 and >8 charged states were rejected and peptide match was disable.

**Label-free quantitative phosphoproteome analyses.** Amastigote (ama) and promastigote (pro) parasites were recovered and washed three times by centrifugation in cold M199 at respectively 2,000 x *g* or 1,600 x *g* for 10 min at 4°C. Samples were incubated for 10 min at 4°C in lysis buffer (1ml per 1.5x$10^9$ promastigotes) consisting of 8 M urea, 50 mM Tris, supplemented with a protease inhibitor cocktail (cOmplete from Roche) and a phosphatase inhibitor cocktail (PhosStop from Roche). Following sonication for 5 min using a sequence of 10 s pulse and 20 s pause, the lysates were centrifuged 15 min at 14,000 x *g* and 4°C, and the supernatant was collected and stored at -80°C until use. Proteins were quantified by RC DC protein assay (Bio-Rad) and adjusted to 1.3 $\mu g.\mu l^{-1}$ in lysis buffer. For detailed protocol see supplementary data file. Disulfide bridges were reduced in 5 mM DTT (Sigma - 43815) for 30 min and alkylated in 20 mM iodoacetamide (Sigma - I1149) for 30 min at room temperature in the dark. Protein samples were diluted 10-fold in 50 mM Tris-HCl and digested with Sequencing Grade Modified Trypsin (Promega - V5111) at a Protein:Trypsin ratio 50:1 overnight. Then a second digestion was performed to complete this step. Proteolysis was stopped by adding formic acid (FA, Fluka - 94318) at a 1% final concentration. Resulting peptides were desalted using Sep-Pak SPE cartridge (Waters) according to the manufacturer's instructions. Peptides were concentrated to almost dryness and were resuspended in 2% ACN / 0.1% FA just before LC-MS/MS injection.

Phosphopeptide enrichment was carried out as described in (2). A slurry of 10 mg/ml of the $TiO_2$ beads (Sachtopore, Sachtleben Chemie, Germany) was prepared in $TiO_2$ buffer (30% ACN / 0.1% trifluoroacetic acid (TFA)). For each sample, GELoader tip was prepared in a Stage-Tips manner using a C8 plug previously activated in methanol and wash with $TiO_2$ buffer. Then, 40 µl of the slurry was packed by centrifugation (100 x *g*) in GELoader tips. The microcolumns were conditioned with 50 µl of loading buffer (80% ACN, 6% TFA, 40 mg/ml glycolic acid) by centrifugation (150 x *g*). The samples were resuspended in loading buffer at a concentration of 2.5 µg/µl and 250 µg were loaded on the microcolumns by centrifugation (100 x *g*). The microcolumns were washed with 50 µl of 80% ACN / 6% TFA and then 100 µl of 50% ACN / 0.1% TFA by centrifugation (150 x *g*). Phosphopeptides were eluted into 60 µl of 10% $NH_4OH$ and then into 5 µl of 80% ACN / 2% FA by centrifugation (100 x *g*). 3.5 µl of 100% FA was added for further acidification.

All analyses were performed on a Q Exactive HF Mass Spectrometer (Thermo Fisher Scientific) coupled with a Proxeon EASY-nLC 1000 (Thermo Fisher Scientific). Phosphopeptides were injected into a home-made 50 cm C18 column (1.9 µm particles, 100 Å pore size, ReproSil-Pur Basic C18, Dr. Maisch GmbH, Ammerbuch-Entringen, Germany). Column equilibration and peptide loading were performed in 250 nl/min in buffer A (0.1% FA). Phosphopeptides were separated at a flow rate of 250 nL/min over 202 min with a multi-step gradient of 2 to 10% buffer B (80% ACN, 0.1% FA) for 40 min, 10 to 30% buffer B for 110 min, 30 to 60% buffer B for 20 min, 60 to 80% buffer B for 1 min. Column temperature was set to 60°C. MS data were acquired using Xcalibur software using a data-dependent method. MS scans were acquired at a resolution of 60,000 and MS/MS scans (fixed first mass 100 m/z) at a resolution of 15,000. The AGC target and maximum injection time for the survey scans and the MS/MS scans were set to 3 x $10^6$ for 20 ms and $10^6$ for 60 ms, respectively. An automatic selection of the 10 most intense precursor ions was activated (Top 10) with a 40 s dynamic exclusion. The isolation window was set to 1.6 m/z and normalized collision energy fixed to 27 for HCD fragmentation. We used an underfill ratio of 1% corresponding to an intensity threshold of $1.7E^5$. No charged states were rejected and peptide match was preferred.

**Proteomics and phosphoproteomics analyses***:* Raw data were analyzed using MaxQuant software (version 1.5.3.8 ) (3) using the Andromeda search engine (4). The MS/MS spectra were searched against the Ld1S database (https://www.ncbi.nlm.nih.gov/bioproject/PRJNA396645, GCA_002243465.1). The settings for the search included (i) trypsin digestion with a maximum of two missed cleavages, (ii) variable modifications for methionine oxidation, N-terminal acetylation and lysine ubiquitinylation, and

(iii) fixed modification for cysteine carbamidomethylation. For phosphorylation events investigation, serine, threonine and tyrosine phosphorylation were added as variable modifications. The minimum peptide length was set to 7 amino acids and the false discovery rate (FDR) for peptide and protein identification was set to 0.01. The main search peptide tolerance was set to 4.5 ppm and to 20 ppm for the MS/MS match tolerance. The setting 'second peptides' was enabled to identify co-fragmentation events. Quantification was performed using the XIC-based Label-free quantification (LFQ) algorithm with the Fast LFQ mode as previously described (5). Unique and razor peptides, including modified peptides, with at least two ratio counts were accepted for quantification.

The workflows applied for the proteomics and phosphoproteomics data analysis are comparable to those previously used in this context (6).

For the differential analyses of proteome data, proteins categorized as 'reverse', 'contaminant' and 'only identified by site' were discarded from the list of identified proteins. After log2 transformation, LFQ values were normalized by median centering within conditions (*normalizeD* function of the R package *DAPAR* (7). Remaining proteins without any LFQ value in one of the conditions (either ama or pro) and at least two values in the other condition were considered as exclusively expressed proteins. Missing values across the four biological replicates were imputed using the imp.norm function of the R package norm (norm: Analysis of multivariate normal datasets with missing values. 2013 R package (version 1.0-9.5)). A limma t-test was applied to determine proteins with a significant difference in abundance while imposing a minimal fold change of 2 between the conditions to conclude that they are differentially abundant (8, 9). An adaptive Benjamini-Hochberg procedure was applied on the resulting p-values using the function *adjust.p* of R package *cp4p* (10) and the robust method described in Pounds et al. (11) to estimate the proportion of true null hypotheses among the set of statistical tests. The proteins associated to an adjusted p-value inferior to a False Discovery Rate (FDR) of 0.01 have been considered as significant and differentially abundant proteins. The volcano-plot is in Figure 2B and contain on their side the iBAQ (12) values of proteins present or absent in one of the two conditions compared.

The workflow of the phosphoproteome analysis is close to the proteomics data analysis with some minor adaptations (13). Phosphosites were only considered if detected with high confidence (identification FDR<1%) and high localisation confidence (localisation probability >0.75) in at least one replicate per condition. This criterion was chosen to retain biologically relevant, low-abundance phosphosites, which are more difficult to identify and are often stochastically sampled in phosphoproteomics datasets. Intensities of proteins and phosphopeptides were normalized by condition using a median-centering function from DAPAR (7), and their missing values were imputed using the impute.mle function of the imp4p R package (14) when at least one observed value was present in that condition. This algorithm imputes values in a condition only when an intensity value has been quantified in at least one of the samples of the considered condition. The quantification profiles (modified peptide and parent unmodified protein quantified/not quantified in at least one sample of a condition) can be viewed in the "Absent/Present" columns of Table 10, sheet A. For the differential analysis of one condition versus another one, two statistical tests were used. First, a moderated t-test was performed thanks to the limma R package (8, 9) to determine whether a phosphorylated peptide is significantly differentially abundant between both conditions. Moreover, phosphorylated peptides quantified in one condition and not in the other were also considered differentially abundant. We applied a contrasted t-test was to compare the variation in abundance of each modified peptide to the one of its parent unmodified protein using the limma R package (8, 9). This second test allows the calculation of a normalized phosphorylation change defined as the ratio of the change in phosphopeptide abundance relative to the change in the corresponding total protein abundance for each condition (paragraph 3.9 in Giai Gianetto et al 2023). An adaptive Benjamini-Hochberg procedure was applied on the resulting p-values thanks to the adjust.p function of R package cp4p (10) using the Pounds et al (11) method to control the False Discovery Rate level. The phosphorylated peptides associated to an adjusted p-value inferior to a FDR of 1% have been considered as significantly and differentially abundant between the compared conditions. Note that this second test can be performed only when there are quantified intensity values for both the non-modified peptide and its parent protein. Interesting cases also emerge from the absence of quantified values for the belonging protein. Consequently, differentially abundant peptides that are associated to proteins from which no fold change can be computed were also considered in the final list of differentially abundant peptides that evolve differently from their parent protein. Results of these differential analyses are summarized in **Figure 5A** and available in Table 10.
